# Multimodal investigation of the neurocognitive deficits underlying dyslexia in adulthood

**DOI:** 10.1101/2024.11.12.623217

**Authors:** Cara Cristina, Zantonello Giulia, Ghio Marta, Tettamanti Marco

**Author notes:** Corresponding author: Marco Tettamanti, Department of Psychology, University of Milan-Bicocca, Milano, Italy Piazza dell’Ateneo Nuovo 1, I-20126 Milano, Italy.

## Abstract

Dyslexia is a neurobiological disorder characterised by reading difficulties, yet its underlying causes remain unclear. Neuroimaging and behavioural studies found anomalous responses in tasks requiring phonological processing, motion perception, and implicit learning, and showed gray and white matter abnormalities in several brain regions of dyslexics compared to controls, indicating that dyslexia is a heterogeneous condition and promoting a multifactorial approach. In order to evaluate whether the combination of behavioural and multimodal MRI can have greater sensitivity in identifying neurocognitive traits of dyslexia compared to monocomponential approaches, in 19 dyslexic and 19 control subjects we acquired behavioural cognitive assessments, multiple (phonological, visual motion, rhythmic) mismatch-response functional MRI tasks, structural diffusion-weighted and T1-weighted images. To examine between-group differences in the multimodal neurocognitive measures, we applied univariate and multivariate approaches. Results showed that dyslexics performed worse than controls in behavioural phonological tasks. Neuroimaging analyses revealed that individuals with dyslexia present reduced cerebellar responses to mismatching rhythmic stimuli, as well as structural disorganization in several white matter tracts and cortical regions previously implicated in dyslexia. Most importantly, in line with the view of dyslexia as a multifactorial phenomenon, a machine learning model trained with features from all three MRI modalities (functional, diffusion, and T1-weighted) discriminated between dyslexics and controls with greater accuracy than models including just one modality. The individual classification scores in the multimodal machine learning model correlated with behavioural reading accuracy. These results confirm that dyslexia should be approached as a composite condition characterised by multiple distinctive cognitive and brain features.

## 1. Introduction

Developmental dyslexia is a specific learning disability characterised by persistent difficulties in reading efficiently despite normal intelligence and adequate educational opportunities (Peterson & Pennington, 2012). Dyslexia represents one of the most common neurobiological disorders, affecting approximately 5-17% of school-aged children (Shaywitz & Shaywitz, 2005). The neurobiology of dyslexia has been widely investigated in the last decades (Norton et al., 2015; Paulesu et al., 2014). Magnetic resonance imaging studies point out gray matter volume (GMV) and cortical thickness (CT) alterations in occipito-temporal, temporo-parietal, and inferior frontal cortices (Ramus et al., 2018), as well as in cerebellar regions (Pernet et al., 2009). Diffusion tensor imaging (DTI) studies further revealed white matter (WM) disorganization in dyslexic subjects affecting association tracts as well as projection and commissural tracts (Cui et al., 2016; Vandermosten, Boets, Wouters, et al., 2012). Functional neuroimaging studies have highlighted the presence of widespread abnormalities in the dyslexic brain, particularly in temporo-parietal, occipito-temporal and inferior frontal cortices (Devoto et al., 2022; Norton et al., 2015; Peterson & Pennington, 2015). The involvement of this extensive network is thought to reflect the multifaceted nature of the cognitive deficits underlying dyslexia (Menghini et al., 2010). Indeed, besides the defining symptomatology of the reading disorder, dyslexic subjects may exhibit impairments in manifold cognitive domains. This evidence has given rise to various theories about the aetiology of dyslexia, with yet little consensus among them (Carioti et al., 2022). Some of the most influential views on the disorder so far stem from the phonological, magnocellular, and cerebellar theories, which posit that dyslexia is caused by deficits in phonological processing (Peterson & Pennington, 2015; Ramus, 2001; Snowling et al., 2020), visuo-motion perception (Bucci et al., 2008; Fischer & Hartnegg, 2000; Gori et al., 2016; Gori & Facoetti, 2015; Stein, 2014, 2019; Tiadi et al., 2016), and implicit learning of rhythmic sequences (Menghini et al., 2006; Vicari et al., 2003), respectively. Although the different theoretical accounts have significantly deepened our understanding of dyslexia aetiology, there is now growing consensus that none of these theories alone can fully account for the aetiology of the disorder. Dyslexia is increasingly recognised as a multifactorial phenomenon, involving a complex interplay of structural, functional brain, and behavioural impairment that collectively contribute to reading difficulties (Peterson & Pennington, 2015). One of the most appropriate methods for examining multidimensionality involves the use of multivariate machine learning (ML) models (Nenning & Langs, 2022; Vu et al., 2018). The recent emergence of ML techniques has yielded substantial benefits in clinical and research contexts (Vogt, 2018; Vu et al., 2018). ML studies have been conducted to predict dyslexia based on various data sources, including psycho-educational tests, web-based games, eye movement tracking, and neuroimaging measurements (Kaisar, 2020; Usman et al., 2021). Specifically, GMV and CT have been used to discriminate between dyslexic and control groups (Płoński et al., 2017; Tamboer et al., 2016) and to predict children’s future literacy skills (Beyer et al., 2022; Tamboer et al., 2016). Similar results were also obtained using WM diffusion indices (Cui et al., 2016; Hoeft et al., 2011; Langer et al., 2017) and functional magnetic resonance imaging (fMRI) data (Hoeft et al., 2011; Vandermosten et al., 2020; Yu et al., 2020; Zahia et al., 2020). However, no study so far has attempted to integrate multimodal brain features into a unique model to predict the diagnosis of dyslexia.

### Aim of the study

Within this framework, the current study aimed to investigate the neurocognitive bases of dyslexia in a group of dyslexic adults compared to a group of matched controls by combining behavioural with functional and structural brain imaging measures. We developed an integrated approach to test the phonological, magnocellular, and cerebellar theories of dyslexia at the cognitive and brain levels within the same experimental paradigm. In addition to behavioural and mismatch response (MMR) fMRI tasks testing the participants’ sensitivity to the phonological, magnocellular, and cerebellar domains, T1-weighted and diffusion-weighted images were also acquired. Between-group differences for each of the cognitive and neuroimaging measures were tested employing univariate and multivariate approaches. Our prediction was that individuals with dyslexia would perform worse than controls in all the cognitive domains and show reduced MMR in all three fMRI tasks. Significant differences between the two groups were also expected for structural gray and white matter indices. Embracing the view of dyslexia as a multifactorial phenomenon, the main aim of the current study was to implement a ML algorithm to evaluate whether the combination of multimodal brain measures could accurately discriminate between subjects with and without dyslexia. We expected the ML model including functional and structural brain measures to outperform both univariate and multivariate models that only include unimodal measures, offering a more comprehensive understanding of the neurocognitive bases of the reading disorder.

## 2. Materials and Methods

Nineteen subjects with a diagnosis of dyslexia (10 males, mean age = 23.67 years, SD = 4.4 years) and 19 control subjects (7 males, mean age = 23.55 years, SD = 2.5 years) participated in the present study. Inclusion criteria for the dyslexic group were: (1) an official diagnosis of dyslexia; (2) text reading speed and/or accuracy score equal to or below the 5^th^ percentile of the normative data in the MT-16-19 test battery (Cornoldi & Candela, 2015); (3) abstract reasoning skills in the normal range. Inclusion criteria for the control group were: (1) no history or familiarity with developmental learning disabilities; (2) reading speed and accuracy scores equal to or above the 10^th^ percentile of the MT-16-19 normative data (Cornoldi & Candela, 2015); (3) abstract reasoning skills in the normal range. Two dyslexic participants did not provide their official diagnosis certificates; however, one was referred by a local dyslexia association, and the other had reading accuracy below the 5^th^ percentile of normative data.

### 2.1 Participants

The two groups were not significantly different for sociodemographic and educational variables (Table 1). Thirteen out of the 19 dyslexic participants reported the presence of comorbidities with other learning disorders (Table 1). All participants were right-handed, Italian native-speakers with no reported history of neurological or psychiatric conditions, brain trauma, substance abuse, or pharmacological treatment affecting the nervous system, and no sensory or perceptual deficits except for corrected-to-normal vision. Participants gave written consent to participate in the study after receiving an explanation of the procedures. The study was approved by the Ethical Committee of the University of Trento, Italy.

**Table 1.** Participants’ demographic information.

|  | **Dyslexics** | **Controls** | **p-value** |
| --- | --- | --- | --- |
| Number of participants | 19 | 19 | - |
| Sex - Male/Female | 10/9 | 7/12 | .5 |
| Age years- mean (SD) | 23.67 (4.40) | 23.55 (2.46) | .91 |
| Education years – mean (SD) | 15 (2.69) | 16.05 (1.68) | .16 |
| Parental education years – mean (SD) | 15.53 (4.12) | 13.39 (2.50) | .06 |
| Comorbidities | Dysgraphia: n = 3<br>Disorthography: n = 6<br>Dyscalculia: n = 10 | - | - |

### 2.2 Experimental design

#### 2.2.1 Cognitive tests

The short version of the Raven’s Advanced Progressive Matrices (RAPM; Arthur et al., 1999; Arthur & Day, 1994) was administered to assess participants’ abstract reasoning and general intelligence. The Italian standardised MT-16-19 reading battery (Cornoldi & Candela, 2015) was used to assess text reading speed and accuracy. To examine the cognitive abilities underlying the three alternative theories of dyslexia, phonological, magnocellular, and cerebellar tests were administered. Phonological abilities were assessed using the standardised VALS Italian test battery (George & Pech-Georgel, 2017), including the following subtests: phonological awareness (PA) segmentation task, PA fusion task, rapid automatised naming (RAN) task, immediate and delayed recall, digit span forward and backward (see Supplementary Methods S1.1, for a detailed description of the phonological subtests). The coherence motion perception (CMP) task was used for testing processing in the magnocellular system (Benassi et al., 2010; Boets et al., 2006; Joo et al., 2017; Kevan & Pammer, 2008). In this task, a kinematogram included a certain proportion of dots moving coherently leftwards or rightwards within the rectangular field, while the remaining dots moved in random directions. Participants were asked to indicate the prevalent direction of the dots’ movement by pressing either the left or the right arrow on a computer keyboard. The individual motion coherence threshold was defined as the smallest fraction of coherently moving dots required for correct direction discrimination (see Supplementary Methods S1.2, for a detailed description of the CMP task). Participants’ implicit learning skills related to cerebellar activity were tested using a serial reaction time (SRT) task (Menghini et al., 2006; Vicari et al., 2003; Yang et al., 2013). Participants were presented with red, blue, and green colored circles (radius = 100 pixels) that appeared sequentially on a computer screen, and they were asked to press the space bar on the computer keyboard whenever they saw a green circle. The entire set of stimuli was organised into five different blocks of 70 stimuli each, with the first block (I) characterised by a pseudo-random stimulus presentation order, followed by three blocks (II-IV) in which a fixed sequence of presentation was implemented, followed by a final block (V) with again pseudo-random order presentation (see Supplementary Methods S1.3, for a detailed description of the SRT task). Reaction times (RTs) and accuracy in response to the green circles were measured. Both the CMP and SRT tasks were created and presented using Psychopy (version 2021.1.2; Peirce et al., 2019). The participants sat at a distance of approximately 60 cm from the screen.

#### 2.2.2 fMRI tasks

We designed three fMRI tasks based on an oddball paradigm. In magneto/electroencephalographic studies, mismatch negativity is an effect elicited by the occurrence of an “oddball” (deviant) stimulus in a repetitive stream of frequent (standard) stimuli, reflecting the individual brain’s sensitivity to perceive and discriminate stimuli (Fitzgerald & Todd, 2020; Garrido et al., 2009; Gu & Bi, 2020; Kujala & N äätänen, 2001; Näätänen et al., 2007). Several studies provided evidence that fMRI captures an equivalent MMR both in auditory and visual modalities (Gomot et al., 2006; Kovarski et al., 2021; Schall et al., 2003).

Three distinct experimental MMR fMRI tasks were designed for the current study, each associated with a matched control MMR task: i) a phonological matched to an auditory task; ii) a magnocellular matched to a parvocellular task; and iii) a cerebellar matched to a color task. All MMR tasks were presented using Psychopy 2021.1.2, and conformed to a block design, with four standard blocks alternated with four deviant blocks. The standard block contained only standard stimuli (N = 26), while the deviant block comprised 21 standard stimuli (81%) and 5 deviant stimuli (19%) in a pseudo-randomised order (see Supplementary Methods S2 for further details).

##### Phonological MMR fMRI task

The experimental, phonological MMR task featured the /ba/ and /da/ syllables as the standard and deviant stimuli, respectively, to elicit a phonological contrast (Alonso-Búa et al., 2006; Gu & Bi, 2020). The syllable stimuli were produced by a female Italian native speaker and recorded in an anechoic room with a condenser microphone. The control auditory MMR task featured non-linguistic tone stimuli generated in MATLAB R2020b. The standard and deviant tone stimuli had the same fundamental frequencies as the F1 and F2 formants of, respectively, the /ba/ and /da/ syllables. All stimuli lasted 300 ms and were separated by a 700 ms-long ISI (Figure 1A).

**Figure 1.**
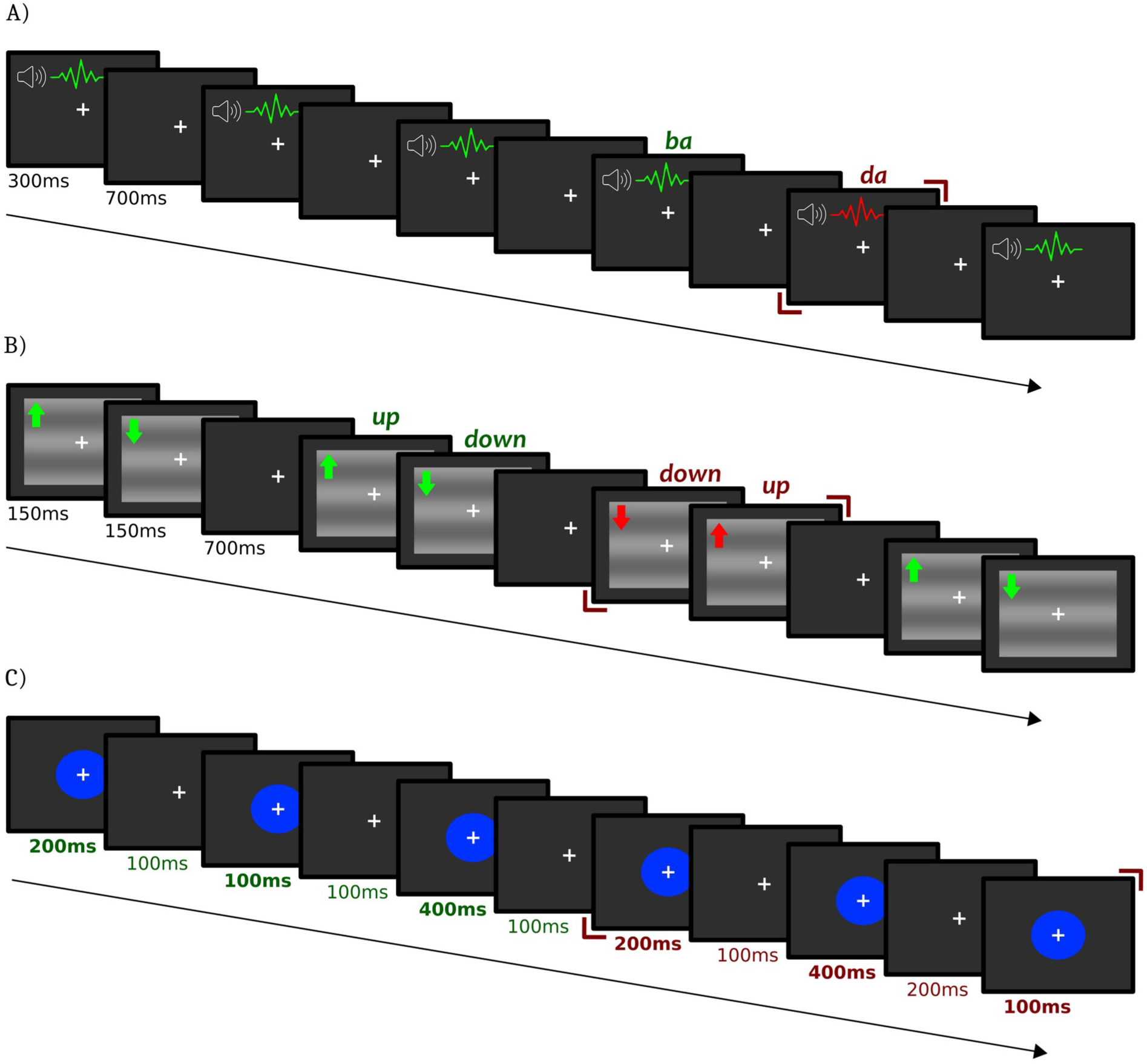
fMRI Mismatch-Response Task Design. Visual representation of standard (green) and deviant (red) stimuli for the phonological, magnocellular and cerebellar MMR tasks. (A) The standard /ba/ and deviant /da/ syllables were used for the phonological task, all stimuli lasting 300 ms with an ISI of 700 ms. (B) The magnocellular task included a standard grating with an upward-downward motion and a deviant grating with a downward-upward motion, all stimuli lasting 300 ms (150 ms in one direction and 150 ms in the opposite direction), with an ISI of 700 ms. For illustration purposes only, the spatial frequency and contrast of the stimuli depicted in this figure have been slightly increased (spatial frequency = 0.2, contrast = 0.2) compared to those used in the actual experiment (spatial frequency = 0.1, contrast = 0.1). (C) The cerebellar task involved a visual rhythmic pattern, with the standard rhythm generated by presenting a blue circle on the screen for 200 – 100 – 400 ms. For the deviant rhythm the second and last stimulus duration were reversed (i.e. 200 – 400 – 100 ms); the ISI lasted for 100ms.

##### Magnocellular MMR fMRI task

The experimental, magnocellular MMR task employed horizontal sinusoidal gratings with spatial frequency of 0.1 cycles per degree, a contrast level of 0.1, and a velocity of 50 degrees per second (Kremláček et al., 2016). For the control parvocellular task, we used horizontal sinusoidal gratings with higher spatial frequency (1 cycle per degree) and contrast levels (1.0), along with a lower temporal frequency (velocity = 5 degrees per second). The standard gratings featured an upward-downward motion pattern (150 ms upward followed by 150 ms downward), while the deviant stimuli displayed a reverse down-up motion sequence (150 ms downward, followed by 150 ms upward). All stimuli measured 10x15 degrees, lasted 300 ms, and were separated by a 700 ms ISI (Figure 1B).

##### Cerebellar MMR fMRI task

The experimental, cerebellar MMR task featured a blue circle (RGB values = 0, 50, 255; radius = 100 pixels) presented in rhythmic sequences. Standard sequences were constructed with a consistent rhythmic pattern featuring stimulus durations of 200 ms, 100 ms, and 400 ms, separated by a constant 100 ms ISI. The deviant sequences were generated by interchanging the second and last stimulus durations within the rhythmic pattern, resulting in durations of 200 ms, 400 ms, and 100 ms, while still maintaining a constant 100 ms ISI. The control color task featured a non-rhythmic presentation of the blue circle used in the experimental task as the standard stimulus, and of a light blue circle (RGB values = 0, 100, 255; radius = 100 pixels) as the deviant stimulus. Both standard and deviant stimuli in the control task were presented for 700 ms, separated by a 300 ms-long ISI (Figure 1C).

### 2.3 Experimental procedure

All participants first underwent the MRI data collection, followed by the behavioural testing session. Participants were provided with MRI-compatible earplugs for listening to the auditory stimuli while being isolated from the scan noise. Every participant underwent six fMRI acquisition runs, with each run including either one of the three experimental MMR tasks or one of the three control tasks. The order of task presentation across the six fMRI runs was counter-balanced across participants, except for the phonological and auditory tasks which were always presented in the first two runs, in order to minimize earplugs displacement during fMRI data acquisition. At the end of the fMRI session, structural T1 images and DWI were acquired.

### 2.4 Neuroimaging data acquisition

MRI scans were acquired at CIMeC, University of Trento, Italy, with a 3-Tesla MRI scanner (MAGNETOM Prisma, Siemens Healthcare, Erlangen, Germany), using a head-neck radio frequency receive coil with 64 channels. Participants first underwent six runs of functional whole-brain image acquisition. Functional images were acquired with a T2*-weighted gradient-echo, simultaneous multislice echoplanar imaging (EPI), and using blood oxygenation-level-dependent (BOLD) contrast (repetition time (TR) = 1000 ms, echo time (TE) = 28.0 ms, 2 mm-isotropic voxels; multi-band acceleration factor = 5). Each run comprised 245 volumes, preceded by 10 dummy scans excluded from subsequent analysis. Each image volume consisted of 65 contiguous axial slices (2 mm thick) acquired in a bottom-to-top interleaved sequence (field of view = 200 mm, in-plane matrix size = 100 x 100). Each run was followed by the acquisition of two echo-planar spin echo sequences (TR = 4010 ms; TE = 28 ms; 2mm isotropic voxels; echo spacing = 0.57 ms; Bandwidth 2272 Hz/Px) with reverse phase encoding directions (i.e., anterior-posterior and posterior-anterior, respectively), which served for geometrical distortion correction of functional images.

High-resolution structural 3D T1-weighted MEMPRAGE images (4 echoes and one root-mean-square (RMS) image) were subsequently acquired (TR = 3280 ms, TE’s = [2.01, 4.04, 6.07, 8.1 ms], 0.7 mm-isotropic voxels, 288 sagittal slices).

Diffusion-weighted data were acquired using a pulse gradient spin echo sequence with the following parameters: resolution = 2 mm isotropic voxels; TR = 4900 ms; TE = 75 ms; shells = b = 0, b = 1000, b = 2000 s/mm^2^, with 12/32/64 directions, respectively, uniformly distributed on a semisphere; anterior-posterior phase encoding direction. This resulted in a total of 108 volumes with 12 b = 0 volumes equally interspersed in the acquisition series. For distortion correction, another sequence with the same parameters but with a shorter duration and reversed phase encoding direction (i.e., posterior-anterior) was acquired.

### 2.5 Behavioural data analyses

Independent two-sample t-tests were conducted to assess significant differences between dyslexic and control subjects for the following behavioural measures: accuracy scores in the RAPM; number of reading errors and reading speed in the Italian MT-16-19 reading battery; accuracy scores for all the phonological subtests of the VALS battery, time required to perform the PA segmentation, PA fusion, and RAN tasks; CMP threshold; and SRT RTs. Significant alpha threshold was set at .05, and Bonferroni multiple testing correction was applied when needed.

### 2.6 Neuroimaging data preprocessing and analyses

#### 2.6.1 fMRI data

##### Preprocessing

Functional brain image pre-processing was performed using SPM12 version 7771 (https://www.fil.ion.ucl.ac.uk/spm/) running on MATLAB R2020b. Structural T1-weighted images were segmented and registered to the Montreal Neurological Institute (MNI) standard space. The preprocessing pipeline for functional images included visual inspection for head motion artefacts (and eventual despiking of artefactual volumes by the mean of neighbouring non-artefactual volumes), spatial realignment, distortion correction, and slice timing. No more than 5% volumes per run were removed for each subject. Functional images were then normalised to the MNI space using the segmentation procedure with the subject-specific segmented structural images as customised segmentation priors. A spatial smoothing with a 4 mm FWHM Gaussian kernel was applied for the univariate fMRI analysis but not for the multivariate analyses.

##### First level analysis

General linear models (GLMs) were specified for each subject with the preprocessed and smoothed functional images, with the time series high-pass filtered at 128 s and globally normalised. We specified three different GLMs. Each model included one experimental task along with the associated control task. Each task (either experimental or control task) was modeled with two regressors of interest (standard or deviant blocks), one nuisance regressor (including task instructions and the distractor blocks (see Methods S2), and the six head movement realignment parameters. Despiked volumes were also modelled as confound regressors. Within each of the three estimated first-level GLMs, we specified the following t-contrasts: 1) the simple MMR effect [deviant blocks > standard blocks] was calculated separately for the experimental task and the control task, by specifying a weight of +1 for the deviant condition and a weight of -1 for the standard condition; 2) the interactions [Task (experimental > control) x MMR (deviant blocks > standard blocks)], testing for stronger MMR in the experimental task compared to the control task (weights: [-1 1 1 -1]; order: [Experimental-standard, Experimental-deviant, Control-standard, Control-deviant]).

##### Second-level region of interest (ROI) analysis

The simple main effect and interaction contrasts specified at the first level were used to specify a set of second-level, random-effects analyses testing for between-group differences in each task. The first-level simple main effect contrast images were used to specify six second-level, two-sample t-test models, investigating whether dyslexic subjects showed altered MMR compared to control subjects in each task: 1) phonological task; 2) auditory task; 3) magnocellular task; 4) parvocellular task; 5) cerebellar task; 6) color task. In turn, the first-level interaction contrast images were used to specify three two-sample t-test models, testing for the mixed three-way interaction between the factors Group (dyslexic > controls), Task (experimental > control), and MMR (deviant blocks > standard blocks) in each task. The significance threshold for these analyses was set at peak-level p < .05, family-wise error (FWE) corrected for multiple comparisons. For each experimental task, we defined a task-specific ROI brain mask that was used to constrain the anatomical space for the second-level analyses, based on previous literature-driven hypotheses (Figure 2A, Supplementary Tables S1A, S1B, S1C).

**Figure 2.**
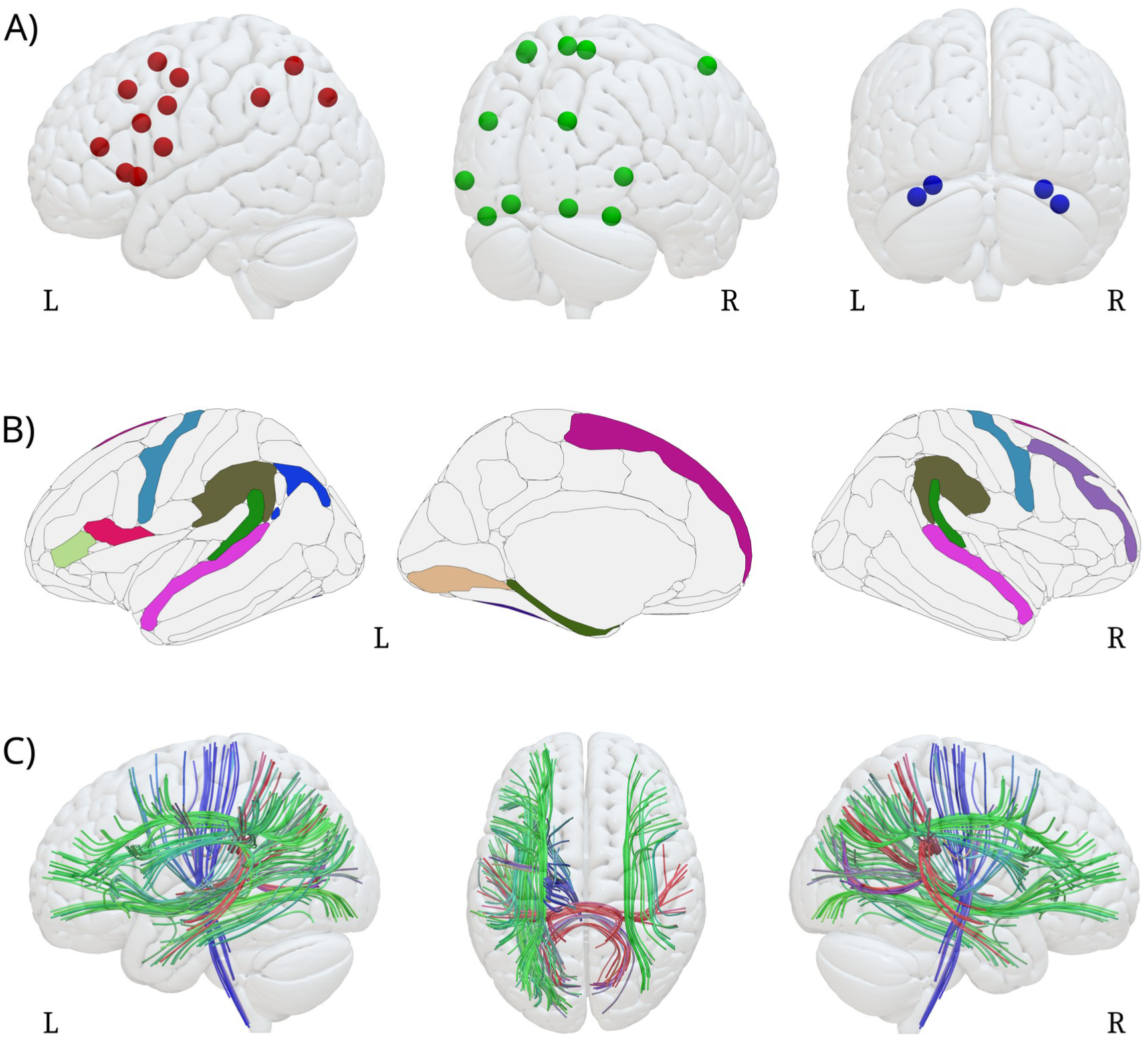
Regions of interest. (A) Phonological (red), magnocellular (green), and cerebellar (blue) task-specific ROI masks used in the second-level, univariate and multivariate fMRI analyses. (B) Regions of interest of the Destrieux Atlas (Destrieux et al., 2010) included in the cortical mask for GMV and CT univariate and multivariate analyses. The different colors identify each a single ROI. (C) Tracts of interest of the White Matter atlas (Radwan et al., 2022) included in the mask used for Fractional Anisotropy and Mean Diffusivity univariate analyses. Red (left-right), green (antero-posterior), and blue (ventro-dorsal) colors represent the prevalent spatial orientation of the individual tracts. ROIs, Regions of interest; GMV, Gray Matter Volume; CT, Cortical Thickness; L, Left; R, Right.

#### 2.6.2 T1-weighted image

##### Preprocessing

T1-weighted images were preprocessed and analysed using FreeSurfer, version 7.2.0 (Fischl, 2012). The main steps of this process included: intensity normalization, registration to Talairach atlas, skull-stripping, white matter segmentation, tessellation of boundaries between gray and WM, deformation to pial surface, automatic topological defects correction (Dale et al., 1999; Fischl et al., 1999). Following preprocessing, the resultant surfaces were visually inspected to ensure accurate segmentation. CT and GMV measures were computed at each vertex using the standard FreeSurfer procedure (Fischl & Dale, 2000; Goto et al., 2022). For univariate analyses only, the individual maps of CT and GMV were smoothed using a full-width half maximum (FWHM) of 10 mm. In addition to the surface-based measures, FreeSurfer also provided segmentation of subcortical regions by assigning labels to each voxel based on probabilistic information. Volumetric measures of the left and right cerebellar cortex were extracted (Fischl et al., 2002).

##### CT and GMV ROI analysis

Alterations in CT and GMV related to dyslexia diagnosis were investigated with vertex-wise two-sample t-tests comparing dyslexic and control subjects, with the total intracranial volume added as a nuisance covariate. Analyses were conducted by fitting a GLM using the FreeSurfer “mri_glmfit” tool. The resulting maps were corrected for multiple comparisons using cluster-based MC simulation with 10000 iterations of randomly generated z-score maps (cluster-wise alpha = .05) (Hagler et al., 2006). To constrain the anatomical space for the analyses of CT and GMV, we defined 20 ROIs (Figure 2B; Supplementary Table S2) based on previous literature (Krafnick et al., 2014; Ma et al., 2015; Ramus et al., 2018; Williams et al., 2018).

#### 2.6.3 DWI data

##### Preprocessing

The preprocessing of DWI data was performed using the Functional Magnetic Resonance Imaging of the Brain (FMRIB) Software Library (FSL 6.0.2; Smith et al., 2004). After visual inspection for motion artefacts, images were corrected for the effects of eddy currents (Horsfield, 1999) and then segmented (non-brain tissue voxels were removed from the whole head images) by the Brain Extraction Tool (Smith, 2002). Diffusion tensor models were then fitted at each voxel with DTIFIT in FDT. This procedure allows to derive the tensor eigenvalues that describe the diffusion coefficients in the primary, secondary, and tertiary diffusion directions, leading to the estimation of fractional anisotropy (FA) and mean diffusivity (MD; Pierpaoli & Basser, 1996).

##### Tract-based spatial statistics ROI analyses

Individual FA images were analysed using Tract-based spatial statistics (TBSS) in FSL (Smith et al., 2006). First, images were aligned to a standard FA template (“FMRIB58_FA”) using a nonlinear registration algorithm. The resulting normalised FA images were averaged to create a mean FA image which was used to generate a mean FA skeleton, representing the centers of all tracts common to all subjects. A threshold of FA > .25 was applied for creating the mean FA skeleton to exclude non-WM voxels. All individual FA and MD maps were then projected onto the mean FA skeleton. Tracts of interest were selected based on previous studies showing their involvement in dyslexia aetiology (Vandermosten, Boets, Wouters, et al., 2012; Zhao et al., 2016, 2022) (Figure 2C; Supplementary Table S3). We defined two GLMs to investigate the effect of dyslexia diagnosis on FA and MD measures within the skeletonised masked images. We then applied the FSL randomise permutation-based program, with 5000 permutations. The significance threshold was set at p < .05 corrected for multiple comparisons by means of threshold-free cluster enhancement (TFCE; Smith & Nichols, 2009).

### 2.7 Correlation analyses

Significant neuroimaging results for the three different MRI modalities (fMRI, GM, WM) were correlated with the behavioural measures that showed significant between-group differences (see paragraph 3.1). In order to avoid a statistical circularity fallacy, the analyses were performed at the regional level, namely if the between-group differences fell within a specific ROI or set of ROIs, the whole ROI(s) - and not only the significant voxels - were used for computing the correlations with the behavioural measures.

### 2.8 Multivariate Pattern Analyses (MVPA)

Multivariate classification between the two groups was performed with the PyMVPA toolbox (version 2.6.5) (www.pymvpa.org; Hanke et al., 2009) running on Python 2.7.18 (www.python.org). The input data for the MVPAs were the same fMRI, GM, and diffusion measures examined at the univariate level.

For each of the three fMRI experimental tasks, one MVPA was performed on the t-statistical images (spmT) derived from the respective simple MMR contrast [deviant blocks > standard blocks] and was anatomically restricted to the task-specific ROI mask (Tables S1A, S1B, S1C). One MVPA model was specified for each of the two GM measures (CT, GMV), by first converting the corresponding surface-based maps to volume-based maps and restricting the analysis to the cortical ROI mask (Table S2). Another MVPA model was defined for each of the two diffusion indices (FA, MD), using the corresponding skeletonised images, and restricting the analysis to the WM tracts of interest (Table S3).

For all MVPAs, we used a C-Support Vector Machine (C-SVM) classifier with a linear kernel (C = -1.0), based on a leave-one-pair-of-subjects-out (LOPSO) cross-validation procedure. In case of a resulting classification accuracy above chance (> 50%), we also performed a searchlight MVPA (Kriegeskorte et al., 2006), with a LOPSO cross-validation procedure and a Gaussian Naive Bayes (GNB) classifier. The searchlight spheres had a radius of 4 mm and were centered at each voxel comprised within the specific mask. The statistical significance was determined with a GNB permutation test (Stelzer et al., 2013).

### 2.9 Multimodal ML model

We implemented a ML model to evaluate whether combining multimodal features could effectively discriminate between subjects with and without dyslexia. The features used in the model corresponded to the fMRI, GM, and diffusion brain measures also analysed with univariate and MVPA approaches. For the three experimental fMRI tasks, GMV, CT, FA, and MD, mean values at the single-subject level were extracted from individual ROI in the respective sets (Tables S1A, S1B, S1C, S2, S3). Total brain volume and mean thickness values of the left and right hemispheres were also extracted (Ramus et al., 2018).

All features for all subjects were concatenated into a matrix with 38 rows (number of subjects) and 102 columns (overall number of ROI features), which was used as input for the ML analysis. The model was implemented using the code released by Cui and colleagues (Cui et al., 2016). The linear C-SVM algorithm with the regularization parameter (C) set to 1 was used to discriminate between the two groups. This discrimination task was performed using a nested-leave-one-out approach, consisting of inner and outer loops. The inner loop served for performing feature selection, and the outer loop served to evaluate classification accuracy. The significance of the accuracy result was determined with permutation tests (N=1000). The most discriminative features were defined as those that had been used in the outer loop of all folds. Classification scores were computed as the distance from the hyperplane and used to determine the class label of each subject (Cui et al., 2016). To compare the performance of combined neuroimaging modalities with individual ones, we implemented four different models: i) the multimodal model including features of all modalities (fMRI, GM, WM), and three single-modality models including ii) only fMRI features, iii) only GM features, iiii) only WM features. To investigate whether the ML classification correlated with the participants’ reading skills, the classification scores of the winning model were correlated with the behavioural performance at reading tests.

## 3. Results

### 3.1 Behavioural results

In the RAPM all participants obtained a score above the 10^th^ percentile of the normative data (Arthur et al., 1999), with no significant differences between the two groups (Supplementary Materials: Table S4). As for the MT-16-19 battery, reading skills differed significantly between the two groups both in terms of the number of reading errors (T(1,36) = 4.35, p = .0002) and of the reading time parameter (T(1,36) = 4.37, p = .0002) (Supplementary Materials: Table S4).

As far as the phonological skills were concerned, dyslexic participants performed more poorly than controls in the following tests: PA segmentation time (T(1,36) = 3.35, p = .018); PA fusion time (T(1,36) = 3.23, p = .026); RAN time (T(1,36) = 4.47, p = .0007); digit span forward score (T(1,36) = -3.75, p = .006); digit span backward score (T(1,36) = -3.54, p = .011). Non-significant results emerged from the following tests: PA segmentation and fusion scores, RAN errors, and immediate and delayed recall scores (Figure 3A; Table S4). No significant between-group differences emerged in the CMP and the SRT tasks (Figure 3B, 3C; Table S4).

**Figure 3.**
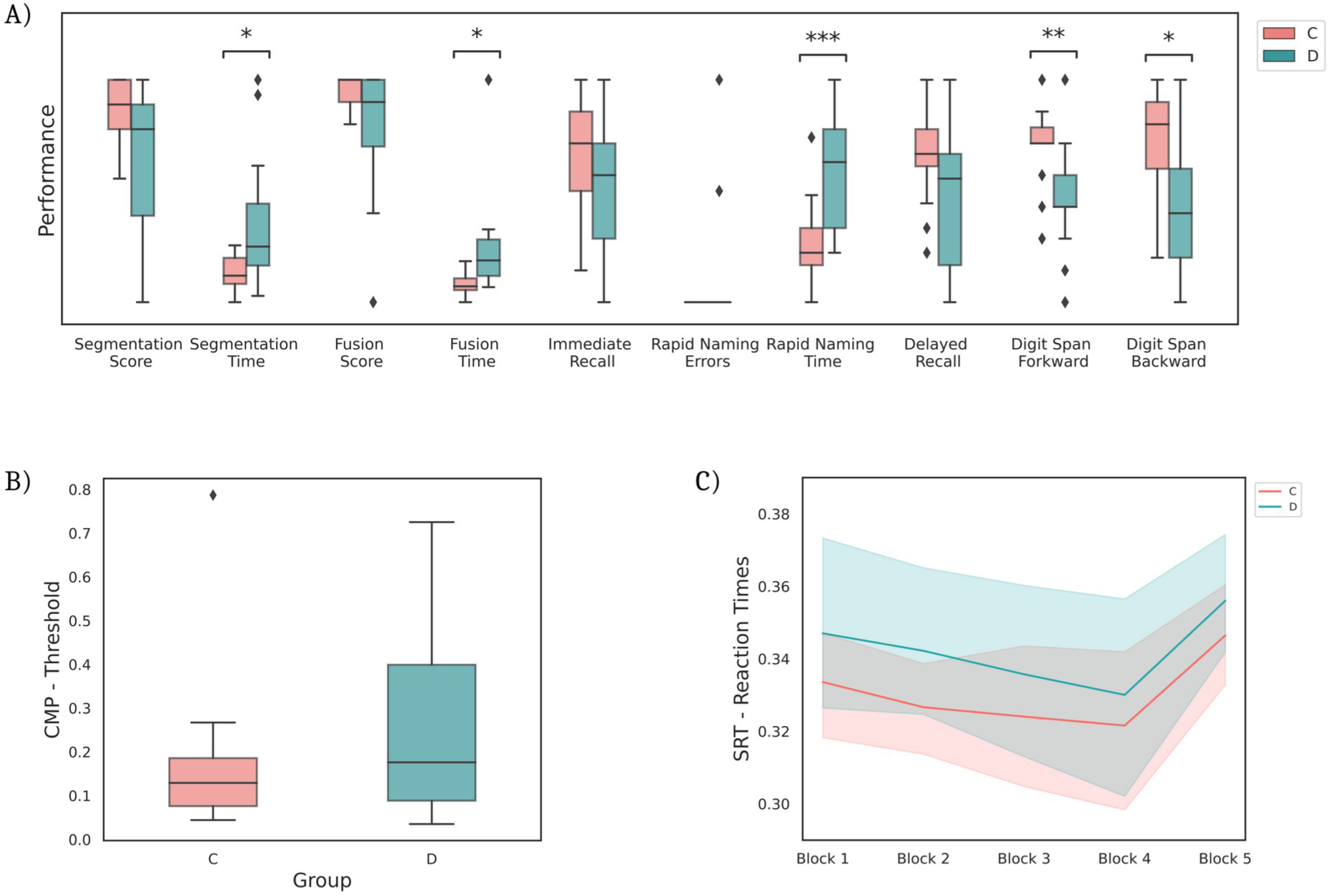
Behavioural results. (A) Significant between groups differences emerged for the following phonological tests: segmentation-time, fusion-time, rapid naming-time, digit span forward, digit span backward. A Bonferroni correction for multiple comparisons (number of tests = 10) was applied. *: p < .05; **: p < .01; ***: p < .001. No significant differences were found for the magnocellular (B) and cerebellar (C) tasks. C, controls; D, dyslexics; CMP, coherent motion perception; SRT, serial reaction time task.

### 3.2 Univariate fMRI results

Regardless of the (dyslexic, control) group, the fMRI tasks elicited detectable MMR activations (Supplementary Table S5).

When testing for between-group (dyslexic vs. controls) differences in each of the experimental tasks, no significant MMR differences emerged in the phonological and the magnocellular tasks. In turn, a significant MMR difference was found in the rhythmic cerebellar task: dyslexic participants showed reduced MMR compared to controls in the right cerebellar lobule VI (peak-level p_FWE_ = .033, Z(1,36) = 3.77, 1 voxel, x = 34, y = -58, z = -26; Figure 4A). Given this significant result, we correlated the cerebellar MMR activation (averaged over all voxels of the anatomical ROI, see Methods 2.7) with the participants’ performance in the behavioural tests. The results showed a significant positive correlation between the digit span forward task and the MMR in the right cerebellum, lobule VI (P_FWE_ = .004, Z(1,36) = 4.33, 5 voxels, x = 36, y = -58, z = -26; Figure 4B).

Results for the control tasks are reported in Supplementary Results S1. Data analyses investigating the interaction effect between the factors Group, Task, and MMR revealed no significant results.

**Figure 4.**
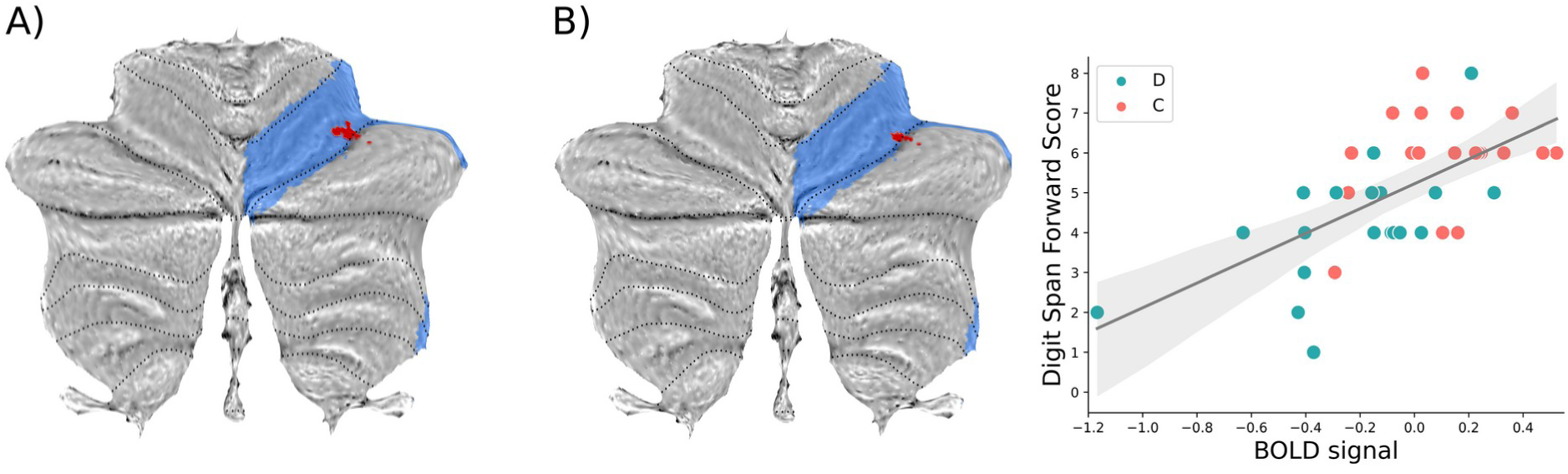
Univariate fMRI results. A) MMR activation in Control > Dyslexic participants in the experimental cerebellar task. Significant voxels (in red; p < .05, FWE-corrected at cluster-level for illustration purposes) are superimposed on the anatomical region corresponding to cerebellar lobule VI (in light blue) extracted from the cerebellar atlas by Diedrichsen et al. (2009). B) Positive correlation between the cerebellar MMR and the digit span forward scores (significant voxels in red, p < .05, peak-level FWE-corrected; cerebellar lobule VI, in light blue) The scatter plot shows the average BOLD signal across significant voxels against the digit span forward scores. BOLD, blood oxygenation level dependent; C, controls; D, dyslexics.

### 3.3 Univariate T1-weighted results

We found no significant between-group differences.

### 3.4 Univariate TBSS results

Between-group FA and MD differences were found in WM tracts of interest (Table S3). More specifically, dyslexic subjects showed reduced FA compared to controls in the third segment of the left superior longitudinal fasciculus (SLF III) (Figure 5A; Table 2A). MD was significantly higher in the dyslexic group in several WM tracts of the left hemisphere, including the SLF, arcuate fasciculus (AF), cortico-spinal tract (CST), thalamo-cortical radation (ThR), middle longitudinal fasciculus (MdLF), inferior longitudinal fasciculus (SLF), inferior fronto-occipital fasciculus (IFOF), and the posterior portions of the corpus callosum (Figure 5B; Table 2A). Based on these results, we correlated FA and MD in the significant WM tracts (see Methods 2.7) with the participants’ performance in the behavioural tests. The results showed significant negative correlations between FA in the SLF III and: i) the number of reading errors; ii) the time required for performing the PA fusion task. Significant positive correlations were found between FA in the SLF III and the time required for performing i) the phonological fusion task, and ii) the PA segmentation task (Figure 5C; Table 2B). MD across a wide set of tracts, including the left AF, SLF, CST and ThR, was positively correlated with the number of reading errors. In turn, a negative correlation was found between the accuracy in the Digit Span Backward task and MD in several tracts, including the left SLF, AF, ThR, IFOF and the posterior part of the corpus callosum (Figure 5D; Table 2B).

**Table 2.**
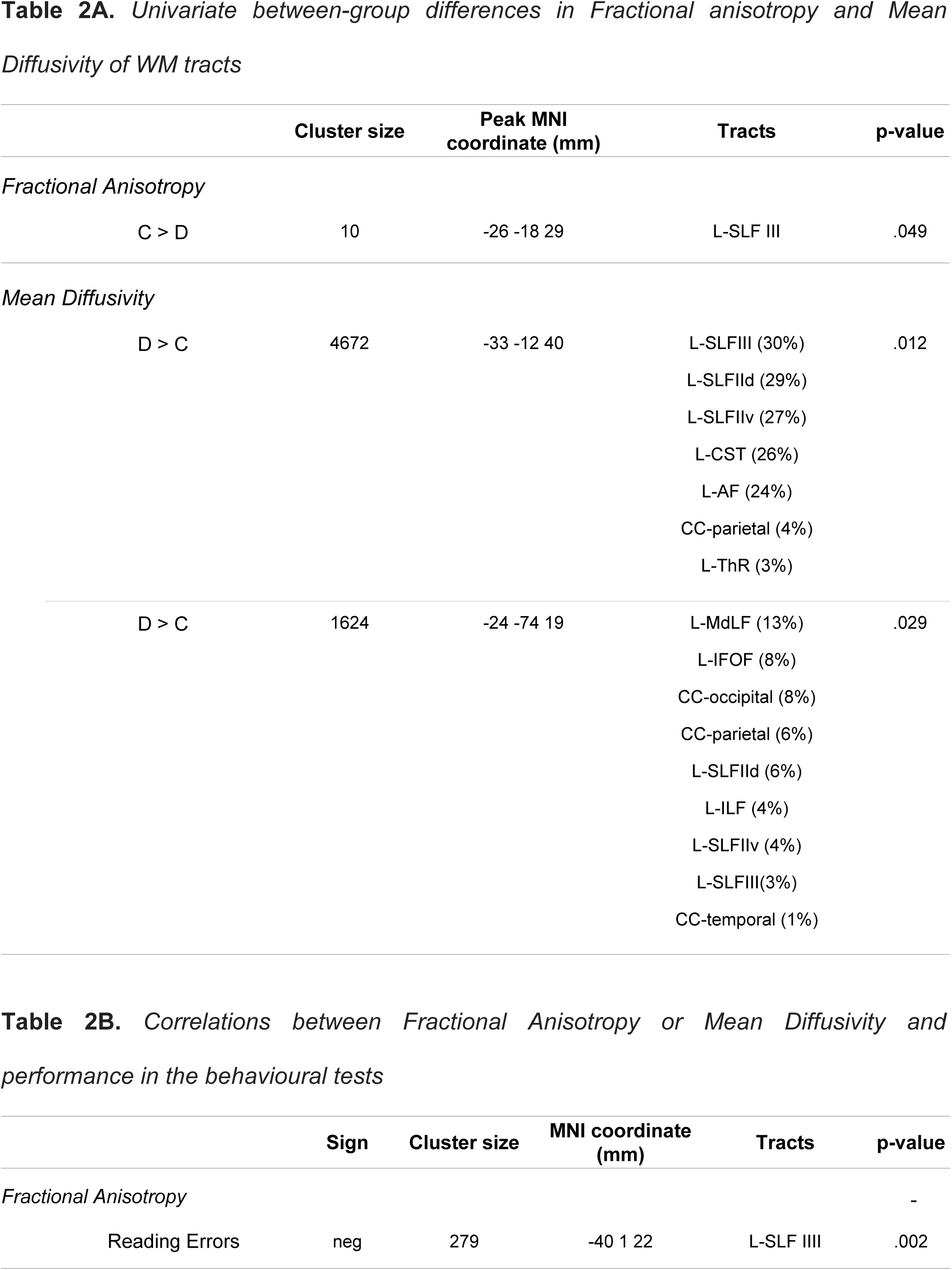
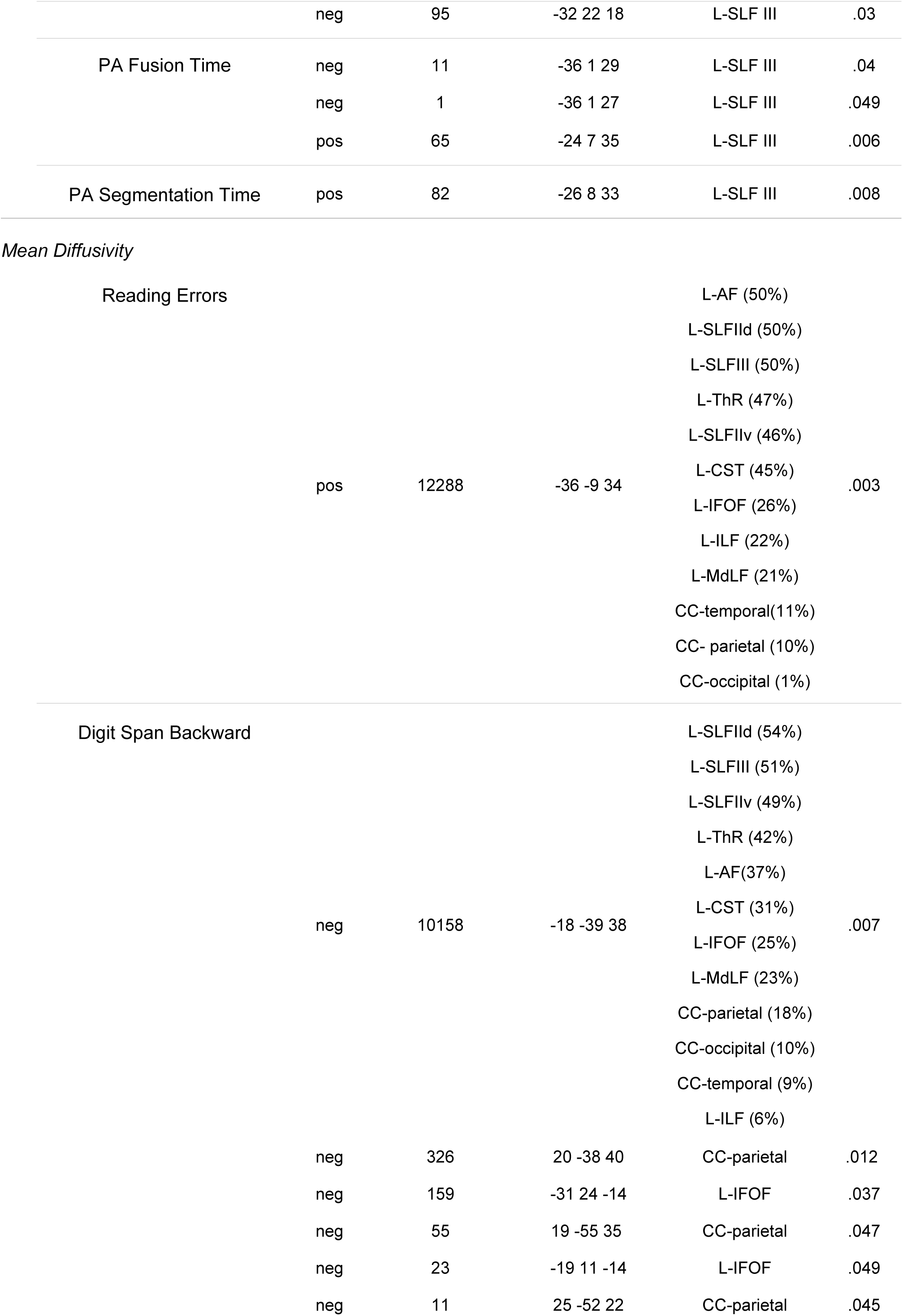
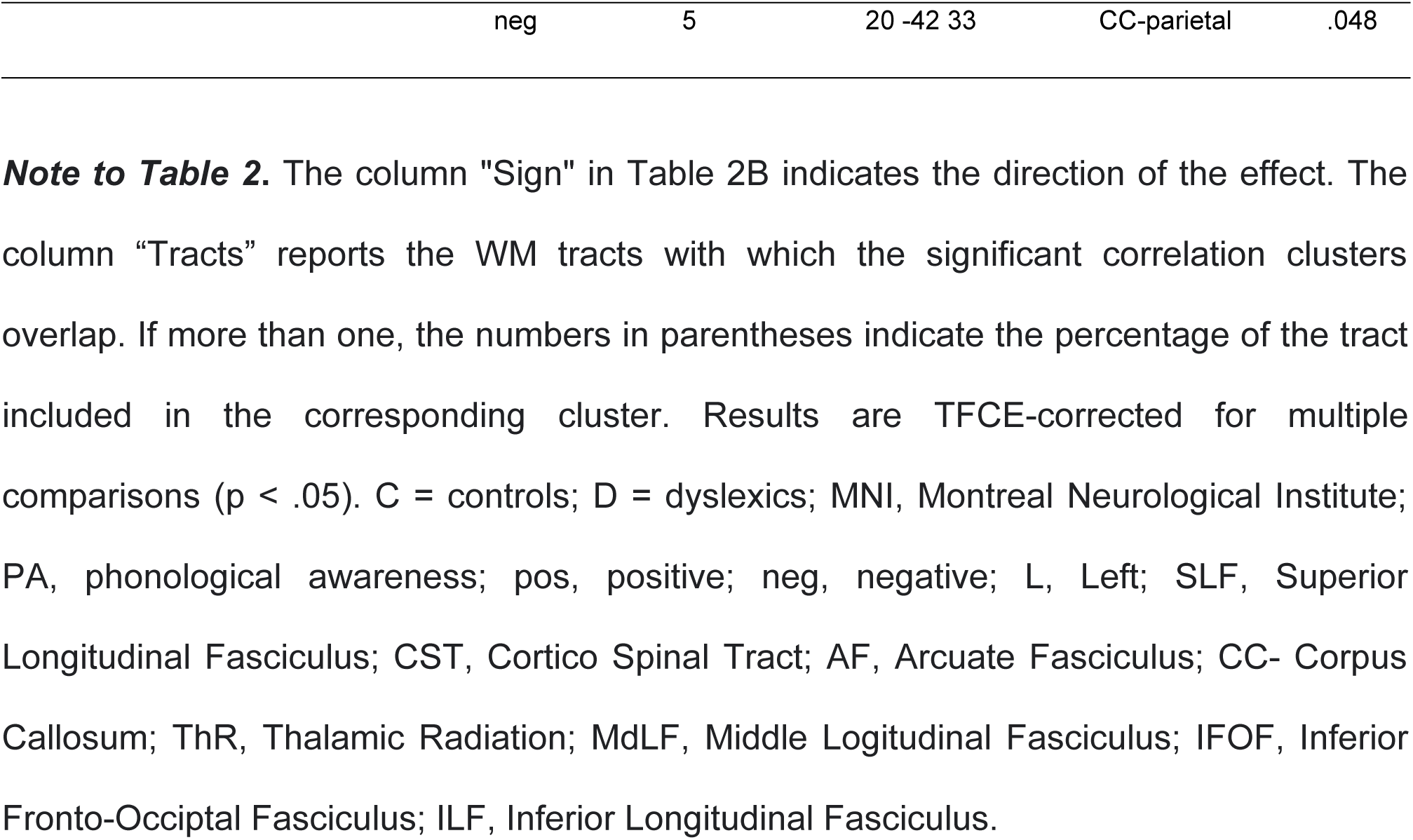
Univariate Fractional anisotropy and mean diffusivity results.

**Figure 5.**
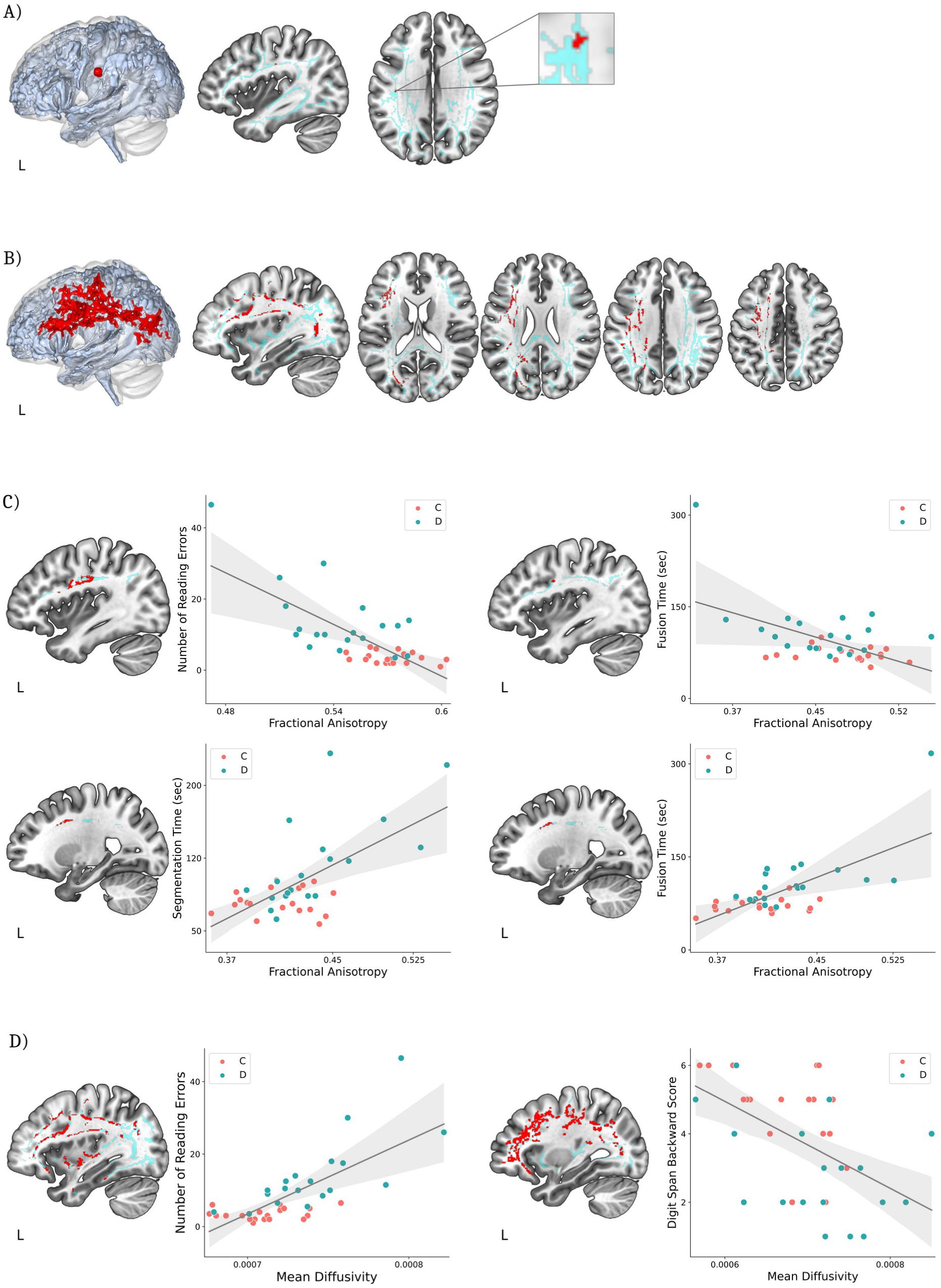
Univariate Fractional anisotropy and mean diffusivity results. Compared to controls, dyslexic participants showed (A) reduced FA in the superior longitudinal fasciculus (SLFIII) and (B) increased MD in several white matter tracts of the left hemisphere. The significant voxels (p < .05, TFCE-corrected) are superimposed (in red) on the TBSS skeleton masked with the a-priori selected WM ROIs (in light blue). (C) Correlations between FA and behavioural tests. The significant voxels (p < .05, TFCE-corrected) are superimposed (in red) on the TBSS skeleton masked with the SLFIII tract (in light blue). The scatter plots show FA averages in the significant clusters (x-axis) against behavioural performances (y-axis). (D) Correlations between MD and behavioural tests. The significant voxels (p < .05, TFCE-corrected) are superimposed (in red) on the TBSS skeleton masked with the WM tracts showing significant MD differences between groups (in light blue). The scatter plots show MD averages in the significant cluster (x-axis) against behavioural performances (y-axis). C, Controls; D, Dyslexics; L, Left.; FA, Fractional Anisotropy; MD, Mean Diffusivity; WM, White Matter; ROIs, Regions of interest; TBSS, Tract based spatial statistics; sec, seconds.

### 3.5 MVPA results

The mean classification accuracy of the MVPA classifier was below the chance level (50%) for both the phonological (48.8%) and magnocellular (40.16%) tasks. In turn, the mean accuracy for the cerebellar task was 62.46% (Figure 6A). The searchlight analysis on the cerebellar task indicated that the sphere yielding the most accurate between-group classification accuracy (74.79%; Figure 6B) was located in the right cerebellar lobule VI (p = .0014 against 20500 permutations, not passing Bonferroni correction for the number of spheres = 1007; x = 34, y = -48, z = -28; Figure 6C).

**Figure 6.**
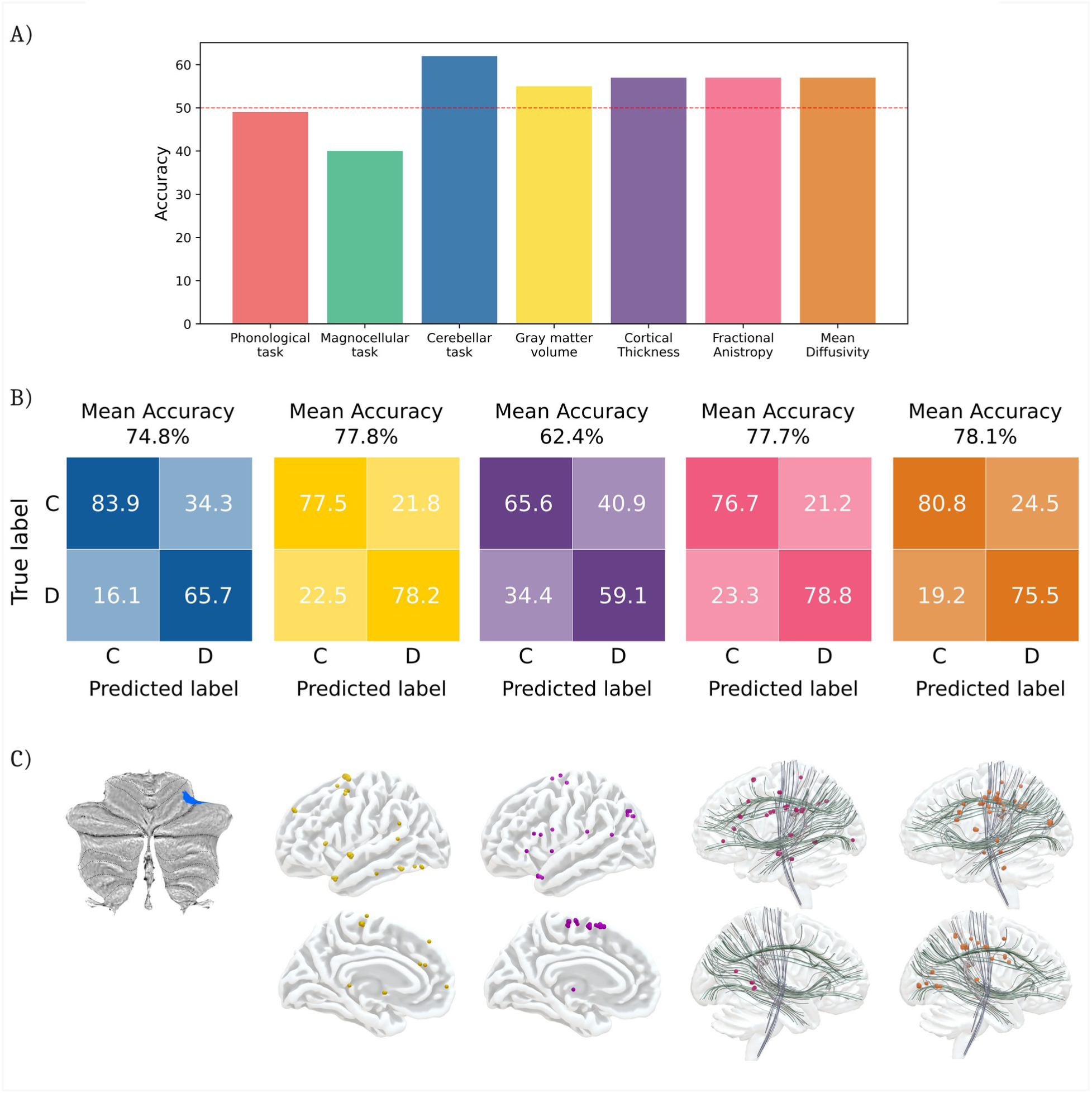
Multivariate pattern analyses results. A) Mean accuracy of the MVPA classifier trained with features from the following modalities: the experimental phonological task (red), experimental magnocellular task (green), experimental cerebellar task (blue), gray matter volume (yellow), cortical thickness (purple), fractional anisotropy (pink), mean diffusivity (orange). The red dotted line represents the chance-level accuracy of 50%. B) Confusion matrices displaying the percent classification accuracies across all significant (p < .0001) searchlight spheres. C) Visual representation of the significant (p < .0001) searchlight spheres for each modality (see Supplementary Table S6). The colors in panels (B) and (C) correspond to those used in (A). C, Controls; D, Dyslexics.

As for GM measures, the MVPA classifier reached a mean classification accuracy of 55% for GMV and 57% for CT (Figure 6A). The searchlight analysis on GMV revealed 80 searchlight spheres with a mean between-group classification accuracy of 77.8% (Figure 6B), which were located in left inferior frontal and occipito-temporal gyri, right middle frontal and precentral cortices, and bilateral superior frontal and temporal gyri (p < .0001 against 10000 permutations; Bonferroni correction could not be assessed due to the exceedingly high computational load for the number of tested spheres = 80928; Figure 6C, Supplementary Table S6). The searchlight analysis on CT yielded 68 significant spheres with a mean between-group classification accuracy of 62.4% (Figure 6B), which were located in the left inferior frontal gyrus, left parietal cortices, bilateral superior frontal, precentral and superior temporal gyri (p < .0001 against 10000 permutations, no Bonferroni correction, number of spheres = 80928; Figure 6C, Supplementary Table S6).

As for WM diffusion indices, the mean classification accuracy was 57% for both FA and MD (Figure 6A). The searchlight analysis on FA revealed 62 spheres with mean between-group classification accuracy of 77.7% (Figure 6B), overlapping with all WM tracts of interest (p < .0001 against 10000 permutations, no Bonferroni correction, number of spheres = 53863; Figure 6C, Supplementary Table S6). The searchlight analysis on MD yielded 147 spheres with a mean accuracy of 78.1% (Figure 6B), overlapping with all WM tracts of interest except the left uncinate fasciculus (p < .0001 against 10000 permutations, no Bonferroni correction, number of spheres = 53863; Figure 6C, Supplementary Table S6).

### 3.6 Machine learning results

The machine learning model including features from all modalities discriminated between the two groups with an accuracy of 66%. The AUC was 0.62, whereas sensitivity and specificity were 68% and 63%, respectively. However, accuracy in this model did not reach significance in the permutation test (p = .1). Compared to the all-modality model, the three single-modality models (fMRI, GM, WM) yielded worse performances with, respectively, an accuracy of 31%, 34%, and 55% (Figure 7).

Within the all-modalities model, the algorithm selected the set of multi-modal features leading to between-group separation. These included three fMRI features, related to MMR in the bilateral cerebellum during the experimental cerebellar task and in the left inferior frontal gyrus during the phonological task; three GM features, including CT in the left occipito-temporal and inferior frontal gyri, and GMV in the planum temporale of the left superior temporal gyrus; and 25 WM features, including both FA and MD measures (Figure 7, Table 3).

By correlating the classification score of each participant in the all-modality model with the behavioural performance at reading tests, we found a significant positive correlation with the number of reading errors (Spearman’s rho = 0.58, p = .0001; Figure 7).

**Table 3.** Discriminant features of the all-modality ML model.

|  | **Modality** | **Measures** | **Features** |
| --- | --- | --- | --- |
| 1 | fMRI | cerebellar-rhythm task | Left Cerebellum (lobule VI) |
| 2 | fMRI | phonological-syllables task | Left Inferior Frontal Gyrus - pars triangularis |
| 3 | fMRI | cerebellar-rhythm task | Right Cerebellum (lobule VI) |
| 4 | GM | CT | L Occipito-temporal Gyrus medial-Parahippocampus |
| 5 | GM | CT | L Inferior Frontal Gyrus - pars triangularis |
| 6 | GM | GMV | L_Superior Temporal Gyrus - planum temporale |
| 7 | WM | FA | L Superior Longitudinal Fasciculus – III |
| 8 | WM | FA | R Superior Longitudinal Fasciculus – II dorsal |
| 9 | WM | FA | L Superior Longitudinal Fasciculus – II ventral |
| 10 | WM | FA | L Superior Longitudinal Fasciculus – II dorsal |
| 11 | WM | FA | L Middle Longitudinal Fasciculus |
| 12 | WM | FA | Corpus Callosum – Parietal |
| 13 | WM | FA | L Inferior Fronto-Occipital Fasciculus |
| 14 | WM | FA | L Uncinate Fasciculus |
| 15 | WM | FA | L Inferior longitudinal Fasciculus |
| 16 | WM | FA | L Arcuate Fasciculus |
| 17 | WM | MD | L Cortico Spinal Tract |
| 18 | WM | MD | R Superior Longitudinal Fasciculus – II ventral |
| 19 | WM | MD | L Arcuate Fasciculus |
| 20 | WM | MD | L Superior Longitudinal Fasciculus – II ventral |
| 21 | WM | MD | R Superior Longitudinal Fasciculus – II dorsal |
| 22 | WM | MD | L Superior Longitudinal Fasciculus – III |
| 23 | WM | MD | Corpus Callosum – Parietal |
| 24 | WM | MD | R Superior Longitudinal Fasciculus – III |
| 25 | WM | MD | L Inferior Fronto-Occipital Fasciculus |
| 26 | WM | MD | L Uncinate Fasciculus |
| 27 | WM | MD | L Superior Longitudinal Fasciculus – II dorsal |
| 28 | WM | MD | L Thalamic Radiation |
| 29 | WM | MD | L Middle Longitudinal Fasciculus |
| 30 | WM | MD | L Inferior longitudinal Fasciculus |
| 31 | WM | MD | Corpus Callosum – Occipital |

*Note to Table 3.* WM, white matter; GM, gray matter; fMRI, functional magnetic resonance imaging; FA, fractional anisotropy; MD, mean diffusivity; CT, cortical thickness; GMV, gray matter volume; L, left; R right.

**Figure 7.**
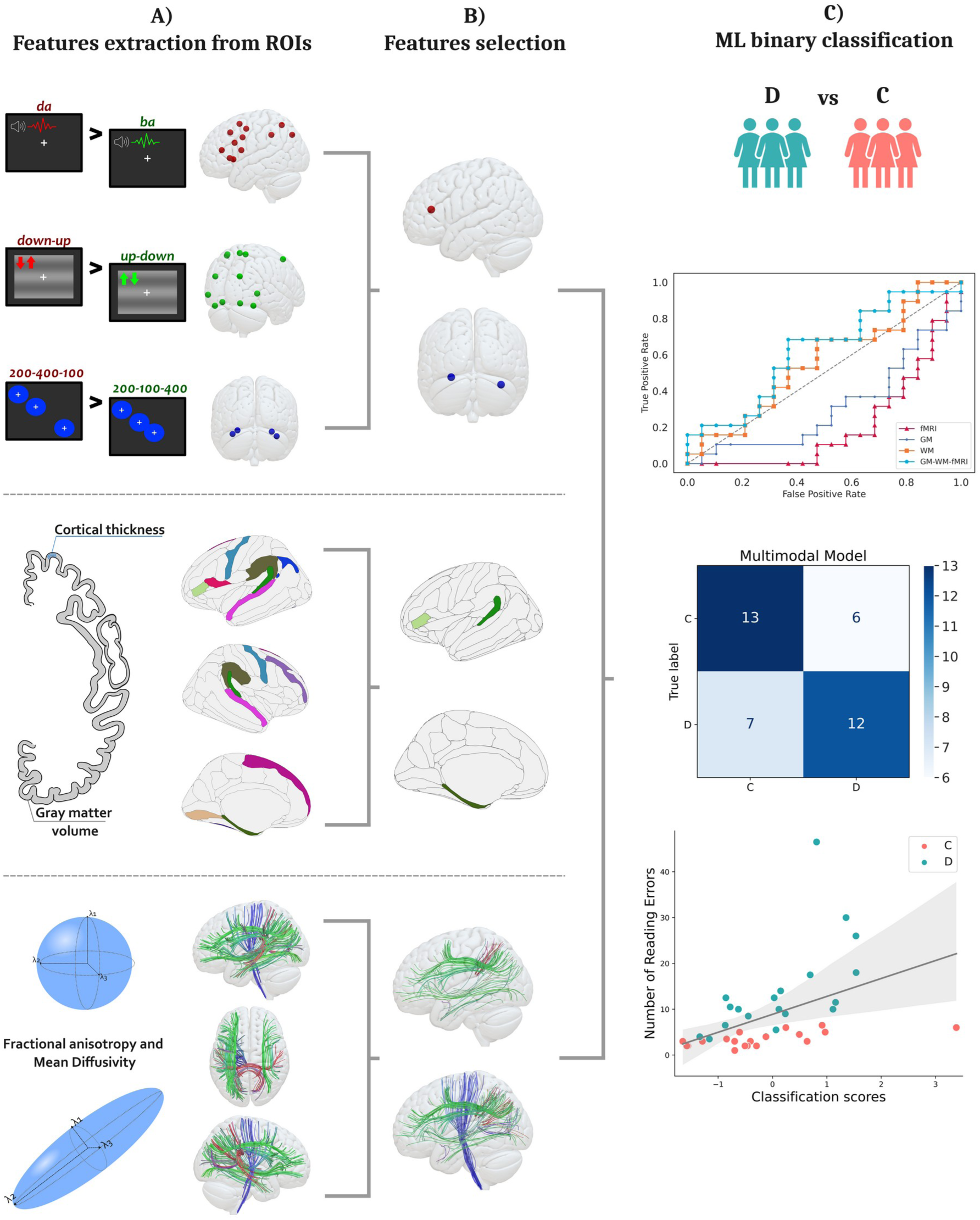
Machine learning model. A) Top: Mean BOLD signal extracted from single-subject data for the three experimental tasks (simple MMR contrast [deviant blocks > standard blocks]) from task-specific regions of interest. Middle: Mean cortical thickness and gray matter volume extracted from anatomical regions of interest. Bottom: Mean fractional anisotropy and mean diffusivity extracted from tracts of interest. B) Discriminative features (i.e., features used in all folds) are shown. Top: BOLD signal in the left inferior frontal gyrus during the phonological task (red) and in the left and right cerebellar lobule VI during the cerebellar task. Middle: Cortical thickness in the left occipito-temporal and inferior frontal gyri and gray matter volume in the left superior temporal gyrus. Bottom: Fractional Anisotropy (top brain) and Mean Diffusivity (bottom brain) in several white matter tracts. C) Top: ROC curve comparison between the multimodal model and single-modality models. Middle: Confusion matrix showing correctly (diagonal) and incorrectly (off-diagonal) classified participants (controls vs. dyslexics). Bottom: Correlation between classification scores (x-axis) and the number of reading errors (y-axis). ROIs, Regions of interest; C, Controls; D, Dyslexic participants; fMRI, functional magnetic resonance imaging; GM, gray matter; WM, white matter.

## 4. Discussion

The current study aimed to investigate dyslexia in adult participants by combining multimodal neuroimaging and behavioural data through an innovative approach. To account for dyslexia heterogeneity, the predictions of some of the most influential theories about the disorder were tested both at the behavioural and brain levels. We first investigated whether dyslexic subjects showed worse behavioural performance in phonological, magnocellular, and cerebellar cognitive tests compared to controls. We then developed a MMR fMRI paradigm to evaluate whether dyslexic subjects showed reduced responses in tasks requiring phonological processing, motion perception, or implicit sequence learning. Structural gray and white matter brain alterations associated with dyslexia diagnosis were also investigated. Between-group differences in each of the examined measures were tested with both univariate and multivariate approaches. As the main aim of the study, we implemented a multimodal machine learning algorithm (Cui et al., 2016) to test whether the combination of multimodal MRI-derived features accurately discriminated between subjects with and without dyslexia.

### Behavioural data show phonological impairment in the dyslexic group

The cognitive assessment revealed that dyslexics were significantly slower but not less accurate than controls in phonological awareness and rapid automatised naming tasks. This aligns with previous literature demonstrating that PA and RAN are universal cognitive markers of dyslexia (Carioti et al., 2022; Devoto et al., 2022), with fluency being a more robust indicator than accuracy (Carioti et al., 2022). These results also support the double-deficit hypothesis, according to which dyslexia arises from distinct deficits in phonological processing and naming speed (Stein, 2023; Vukovic & Siegel, 2006). Dyslexic participants showed worse performance in phonological short-term and working memory skills, assessed through the digit span tasks, consistent with prior research suggesting that working memory training could be a potential treatment for reading and learning disorders (Maehler et al., 2019). In contrast, tests of immediate and delayed recall targeting semantic memory resulted in no significant differences between the two groups, confirming that semantic processing is preserved in dyslexia (Rasamimanana et al., 2020). Unexpectedly, we found no significant between-group differences in the CMP (testing for the magnocellular hypothesis) and SRT (testing for the cerebellar hypothesis) tasks. Despite the lack of statistical significance, the CMP (Figure 3B) and SRT (Figure 3C) response profiles revealed a greater data dispersion in the dyslexic group compared to controls, with a more heterogeneous distribution of data in the upper boundary (i.e., worse performance). This observation may indicate insufficient statistical power to reveal subtle effects, or large within-group variability in our dyslexic sample, with a subgroup of participants with magnocellular or cerebellar impairments, and another subgroup without, in line with the existence of different subtypes of the disorder (Jednoróg et al., 2014; Menghini et al., 2010; Wolf et al., 2024).

### fMRI data show cerebellar impairments in the dyslexic group

Univariate analyses showed reduced MMR in dyslexics compared to controls in the right cerebellar lobule VI during the cerebellar task. MVPA showed that the cerebellar task also provided between-group classification accuracy above chance level. The searchlight analysis revealed that the brain region displaying the highest between-group classification accuracy was located in the right cerebellar lobule VI. This pattern of results is consistent with previous studies showing abnormal activation and reduced gray matter volume in the right cerebellar lobule VI of dyslexic individuals (Menghini et al., 2006; Nicolson et al., 2001; Pernet et al., 2009; Stoodley, 2016). This finding has been linked to the key role of the cerebellum in the predictive brain system (Gatti et al., 2021; Leggio & Molinari, 2015). The cerebellum is known to support the learning of patterns of structured events, and the generation of predictions about the future based on these patterns. It also produces signals when an event deviates from these expectations (Gatti et al., 2021; Leggio & Molinari, 2015). In light of this view, the cerebellum’s involvement aligns with the MMR oddball paradigm, which is thought to reflect the brain’s automatic detection of a prediction error when an incoming stimulus does not match an expected pattern (Fong et al., 2020). In the context of our fMRI study, the cerebellar task was the only one constituted by stimuli intrinsically characterised by a predictable rhythmic structure, thus likely engaging the cerebellar predictive system. Furthermore, the correlation analyses revealed that increased cerebellar activity was associated with better performance at the digit span forward test. This is consistent with the previous literature showing cerebellar involvement in phonological short-term memory (Ashida et al., 2019; Marvel & Desmond, 2010; Peterburs et al., 2019) and outlines a probable connection between cerebellum impairment and phonological deficits (Nicolson et al., 2001; Nicolson & Fawcett, 2019). Against our prediction, we found no significant results for the phonological and magnocellular tasks using either univariate or multivariate models. It remains unclear whether the neurocognitive processes underlying these tasks were compensated to the normal range in our sample of dyslexic adults, or whether there were subtle effects that could not be detected due to insufficient statistical power.

### Structural data show widespread gray and white matter impairments in the dyslexic group

We investigated the effects of dyslexia diagnosis on gray matter volume and cortical thickness of several regions of interest. No significant between-group differences emerged from the univariate analyses. These null results may be linked to the inconsistent evidence about reliable structural markers of dyslexia emerged so far (Ramus et al., 2018). Inconsistencies across studies may be due to several factors, including dyslexia heterogeneity, gender differences, small sample size, and varying methodological approaches (Ramus et al., 2018). However, our multivariate searchlight analyses indicated gray matter measures as potential markers for distinguishing between dyslexic and control groups. More specifically, gray matter volume in frontal, superior temporal, and occipito-temporal regions, along with cortical thickness in the inferior and superior frontal gyri and temporo-parietal regions, showed above-chance between-group classification accuracy. The differing results obtained from univariate and multivariate approaches may suggest that multivariate analyses possess higher sensitivity for detecting subtle and distributed patterns of brain differences. Overall, these results confirm previous evidence that dyslexia is associated with widespread neuroanatomical variations, particularly in regions involved in reading and language processing (Ramus et al., 2018), reinforcing the multifactorial nature of the disorder.

Diffusion tensor imaging univariate analyses revealed widespread disorganization of white matter microstructure associated with dyslexia diagnosis. Between-group comparisons highlighted reduced fractional anisotropy and increased mean diffusivity in several left-hemispheric tracts of dyslexics compared to controls. Consistently, multivariate searchlight analyses showed that diffusion indices in white matter tracts could also discriminate dyslexic from controls subjects with accuracy above chance level. The involvement of these white matter bundles in reading and language processing was corroborated by correlation analyses between diffusion indices and behavioural performance in reading and phonological tests. The results showed that higher fractional anisotropy and reduced mean diffusivity were associated with better reading and phonological performance. These findings align with previous research (Cui et al., 2016; Ramus et al., 2018; Sihvonen et al., 2021; Vandermosten, Boets, Poelmans, et al., 2012; Vandermosten, Boets, Wouters, et al., 2012), providing further support for the idea of dyslexia as a *disconnection syndrome.* According to this perspective, the reading deficit is characterised by disorganised white matter connectivity across distributed brain networks, rather than localised impairments in isolated cortical areas (Habib, 2021; Paulesu et al., 1996; Vandermosten, Boets, Poelmans, et al., 2012).

### Machine learning model

The machine learning model trained with features from all modalities discriminated between dyslexic and control subjects with an accuracy of 66%, though this did not reach significance in the permutation test (p = .1). The single-modality models yielded lower accuracies: 31% for the fMRI model, 34% for the GM model, and 55% for the WM model. The multimodal model showed a significant positive correlation between the classification scores and the number of reading errors. The finding that the multimodal model outperformed single-modality models supports the multifactorial nature of dyslexia, suggesting that different structural and functional brain characteristics may be involved across individuals. Accordingly, the features selected by the all-modality model that mostly contributed to the between-group classification included measures from all three modalities. Consistently with the univariate results, the greatest proportion of selected features was represented by WM measures, including FA and MD in several associative, commissural, and projection tracts. Three fMRI features also contributed to the between-group classification, including MMR in the cerebellar region during the cerebellar task, with good agreement with the univariate and multivariate analyses, but also in the inferior frontal gyrus during the phonological syllable task, in line with previous evidence indicating abnormal activation in the inferior frontal gyrus during phonological tasks in dyslexic subjects compared to controls (Devoto et al., 2022; Georgiewa et al., 2002). The ML model further revealed that cortical thickness in the left occipito-temporal and inferior frontal regions, as well as gray matter volume in the planum temporale of the superior temporal gyrus substantially contributed to between-group classification. This finding aligns with a large body of literature, showing cortical thickness and gray matter volume alterations in occipito-temporal regions of dyslexic subjects (Frye et al., 2010; Kronbichler et al., 2008; Ramus et al., 2018). These regions include the so-called visual word form area, a brain region responsible for letters and word recognition (Caffarra et al., 2021; Kronbichler et al., 2008). As for the inferior frontal gyrus, although no study so far reported a thinner cortex in this region in dyslexic subjects compared to controls, gray matter volume reductions have been previously observed (Brown et al., 2001; Norton et al., 2015; Ramus et al., 2018). Initial evidence of structural abnormalities in the planum temporale of dyslexic brains was provided by Galaburda’s post-mortem studies (Galaburda et al., 1985) and has been largely confirmed by recent evidence (Altarelli et al., 2014; Bloom et al., 2013; Ramus et al., 2018). However, these results should be taken with caution, as it is worth emphasizing that the model did not reach statistical significance in the permutation testing. The classification accuracy of 66% is relatively low compared to similar studies, which have reported accuracies exceeding 80% (Cui et al., 2016; Langer et al., 2017; Tamboer et al., 2016; Zahia et al., 2020). This discrepancy may be due to the limited size of our samples, consistent with the notion that machine learning classifiers take particular advantage of larger datasets for effective training. Nonetheless, the classification scores for the individual participants derived from the all-modality model significantly correlated with individual reading accuracy. This finding is particularly relevant, as it suggests that the combination of multimodal MRI features may offer insights into the individual severity of the reading deficits.

### Limitations and future directions

The current study suffers from some important limitations. First, the relatively small sample size may determine reduced statistical power, potentially hindering the detection of subtle effects and introducing noise in effect estimation. Second, investigating a sample of adult participants with high educational level for studying developmental learning disorders may be sub-optimal, as we cannot exclude compensatory mechanisms developed over time. Moreover, the fact that some dyslexic participants presented comorbidities with other learning disorders may represent a source of confound. Indeed, the cognitive and brain measures here examined are not exclusively related to dyslexia but also show correlations with other learning difficulties (Landerl et al., 2009; Willcutt et al., 2013). Consequently, some of the effects associated with dyslexia may be contaminated by other learning disorders.

Given these limitations, it would be valuable for future research to explore the current paradigm in a paediatric sample. The use of the mismatch negativity is particularly suitable for studying young children, as it captures an implicit and automatic response without requiring active task-related compliance. Furthermore, several studies suggested that phonological mismatch negativity and structural brain measures in preschool children could serve as potential predictors of future reading skills (Kraft et al., 2016; Raschle et al., 2012; Raschle et al., 2011; van Zuijen et al., 2013; Vandermosten et al., 2015). Consequently, longitudinal studies should evaluate whether the integration of multimodal neuroimaging and cognitive measures can predict the outcome of cognitive development and dyslexic impairments. This approach could offer important insights into the early identification and intervention for specific reading disorders.

### Conclusions

Extensive research has shown that dyslexia involves widespread impairments across various cognitive, functional, and structural brain measures. The current study integrates these diverse elements, combining cognitive assessments with functional and structural MRI data to provide a more comprehensive view of the reading disorder. Our findings indicate that our sample of dyslexic adults experience significant phonological difficulties. Functional abnormalities in the right cerebellum, along with widespread disorganization in gray and white matter regions, likely contribute to reading and phonological difficulties. Crucially, this study demonstrated that using a multivariate model to combine diverse neuroimaging measures is the most effective approach to uncovering the complexity of developmental dyslexia.

## Acknowledgements and funding statement

We thank the volunteers who kindly participated in our study. M.G. was supported by a postdoctoral researcher grant by the University of Trento, Italy (Decree No. 48, 23.05.2019) awarded to M.T.

## Data availability statement

The data that support the findings of this study are available on request from the corresponding author. The data are not publicly available due to privacy restrictions.

## Ethics approval statement

The study was approved by the Ethical Committee of the University of Trento, Italy (Protocol no. 2019-014).

## Author contributions

C.C., M.G., M.T. designed the experiment; C.C. and G.Z. collected the data; C.C., G.Z., M.T. analyzed the data; C.C., G.Z., M.G., M.T. organized and wrote the manuscript.

## Supplementary Materials

### Methods S1.1. Phonological battery subtests

- Phonological Awareness (PA) segmentation task: participants were asked to do a letter-by-letter spelling of 10 words spoken aloud by the examiner. Accuracy (number of correctly spelled words) and time (in seconds) required for performing the whole task were computed for each participant.
- PA fusion task: participants had to recognize the words that the examiner spelled aloud letter-by-letter. Accuracy (number of correctly recognized words) and time (in seconds) required for performing the whole task were computed for each participant.
- Rapid Automatized Naming (RAN) task: participants were visually presented with two pages of repeating symbols (i.e., banana, butterfly, kite, airplane), which they were asked to correctly name in succession as quickly as possible. Accuracy (number of correctly named items) and time (in seconds) required for performing the whole task were computed for each participant.
- Immediate and Delayed Recall task: participants were presented with a list of 12 words spoken by the examiner; right after (immediate recall) or after a 3-minute delay (delayed recall), participants were asked to repeat all words, irrespective of the order. The total number of correctly recalled items was computed for each participant, separately for the immediate and delayed recall tests.
- Digit Span Forward task: participants were asked to repeat spans of digits of increasing length in the same order as they were presented. Accuracy was computed for each participant as the number of correctly repeated items.
- Digit Span Backward task: participants were asked to repeat spans of digits of increasing length in the reverse order. Accuracy was computed for each participant as the number of correctly repeated items.

### Methods S1.2. Coherent Motion Perception (CMP) task

A rectangular dot kinematogram of field size 23 x 23 degrees was created comprising 100 white circular dots randomly distributed on a black background (dot diameter = 10 degrees; contrast = 1; speed = 4.2 degrees/sec; dot lifetime = 5 video frames before disappearing and being redrawn in another random location; maximum duration of stimulus presentation = 6 seconds).

The kinematogram included a certain proportion of dots moving coherently leftwards or rightwards within the rectangular field, while the remaining dots moved in random directions. The percentage of motion coherence started at 100% and then decreased by a step size factor of 1.14 (Boets et al., 2006). Participants were asked to indicate the prevalent direction of the dots’ movement by pressing either the left or the right arrow on a computer keyboard. Before starting the actual task, they were presented with six practice trials with high motion coherence (from 0.9 to 1). The motion coherence threshold was defined as the smallest fraction of coherently moving dots required for correct direction discrimination and was estimated with a 2-down-1-up adaptive staircase procedure, the end of which was reached after eight direction reversals (minimum number of trials = 30). The individual threshold was calculated based on the mean of the values in the final four direction reversals.

### Methods S1.3. Serial Reaction Time (SRT) task

Participants were presented with red, blue, and green coloured circles (radius = 100 pixels) that appeared sequentially on a computer screen, and they were asked to press the space bar on the computer keyboard whenever they saw a green circle. The entire set of stimuli was organized into five different blocks of 70 stimuli each, with the first block (I) characterized by a pseudo-random stimulus presentation order, followed by three blocks (II to IV) in which a fixed sequence of presentation was implemented (red-green-blue-red-blue-green-blue), with 10 repetitions in each block, followed by a final block (V) with again pseudo-random order presentation. The pseudo-randomization in blocks I and V was constrained by an equal number of coloured circles as in the fixed-sequence blocks (30 blue, 20 green, 20 red), with no more than two consecutive stimuli with the same colour. Stimulus duration and inter-stimulus interval (ISI) varied randomly from 0.5 to 1 second to avoid the green circles appearing at constant time intervals.

The task was preceded by a familiarization phase during which 18 random stimuli were presented and subjects had to press the space bar at the green circle appearance. Reaction times (RT) and accuracy in response to the green circles were measured. The mean values of the RT in each of the five blocks were calculated for the participants of the two groups and subsequently separately plotted to reveal the groups’ learning curves, representing the increase or decrease of the two groups’ RTs within the task blocks due to the random or sequential stimuli presentation. Similar to the procedure used by Vicari and colleagues (2003), the individual RT differences between blocks IV and I (IV–I), blocks IV and II (IV–II), and blocks V and IV (V–IV) were calculated for both the dyslexic and the control group.

### Methods S2. fMRI tasks

Each fMRI task (either experimental or control) included 188 standard and 20 deviant stimuli, organized in a block-design fashion. For each task, four standard (S) and four deviant (D) blocks were presented in an alternating SDSDSDSD sequence. The standard block contained only standard stimuli (N = 26), while the deviant block comprised 21 standard stimuli (81%) and 5 deviant stimuli (19%). In all tasks, the onset-to-onset interval was set at one second (i.e., 1 Hz frequency). The presentation order of stimuli in the deviant blocks was pseudo-randomised, with the only constraint that each deviant stimulus had to be preceded by a minimum of four standard stimuli. Considering that MMR is an automatic response occurring independently of focused attention (Näätänen et al., 2007), participants were instructed to press a button of an MRI-compatible response box in response to the visual presentation of a sparse distractor stimulus, constituted by a white rectangle. Three distractor blocks were included in each fMRI task, following an SD-distractor-SD-distractor-SD-distractor-SD order. During the distractor blocks, the standard stimulus was presented at the usual 1 Hz frequency for 8 seconds, with the first 4 seconds also featuring the simultaneous presentation of the white rectangle. All stimuli were presented on a dark-gray background (RGB = 46, 46, 46) with a white fixation cross in the center of the screen. In all tasks, stimuli presentation and response collection were controlled using Psychopy software (version 2021.1.2; Peirce et al., 2019).

### Results S1. Between-group MMR differences in the fMRI control tasks

No significant between-group differences emerged for the parvocellular task and for the color task. Dyslexic subjects showed increased MMR in the auditory tone task compared to control subjects in the left superior medial frontal gyrus (peak-level p_FWE_ = .049, Z(1,36) = 3.89, 1 voxel, x= -4, y= 16, z= 42).

---

**Table S1.**
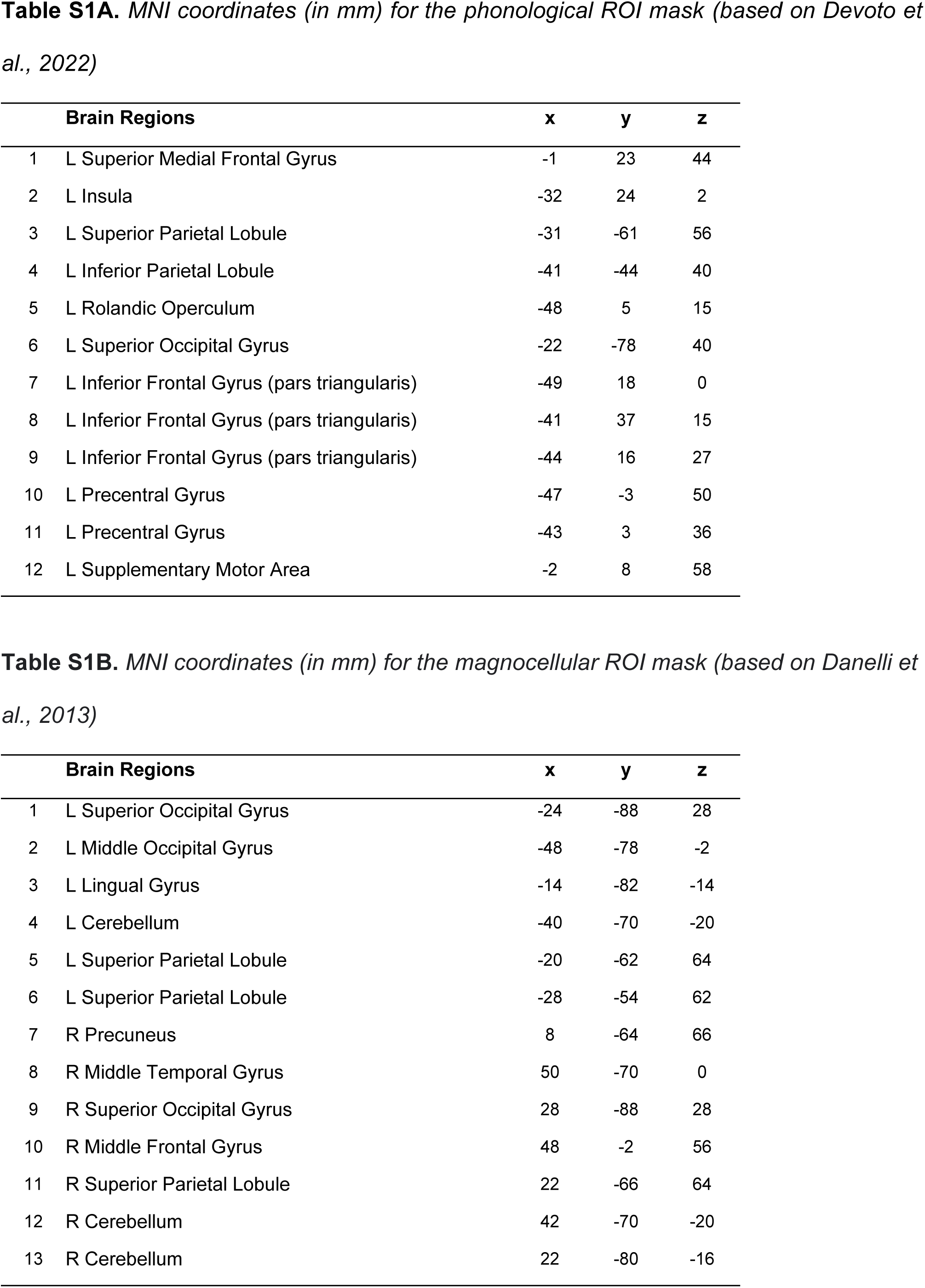
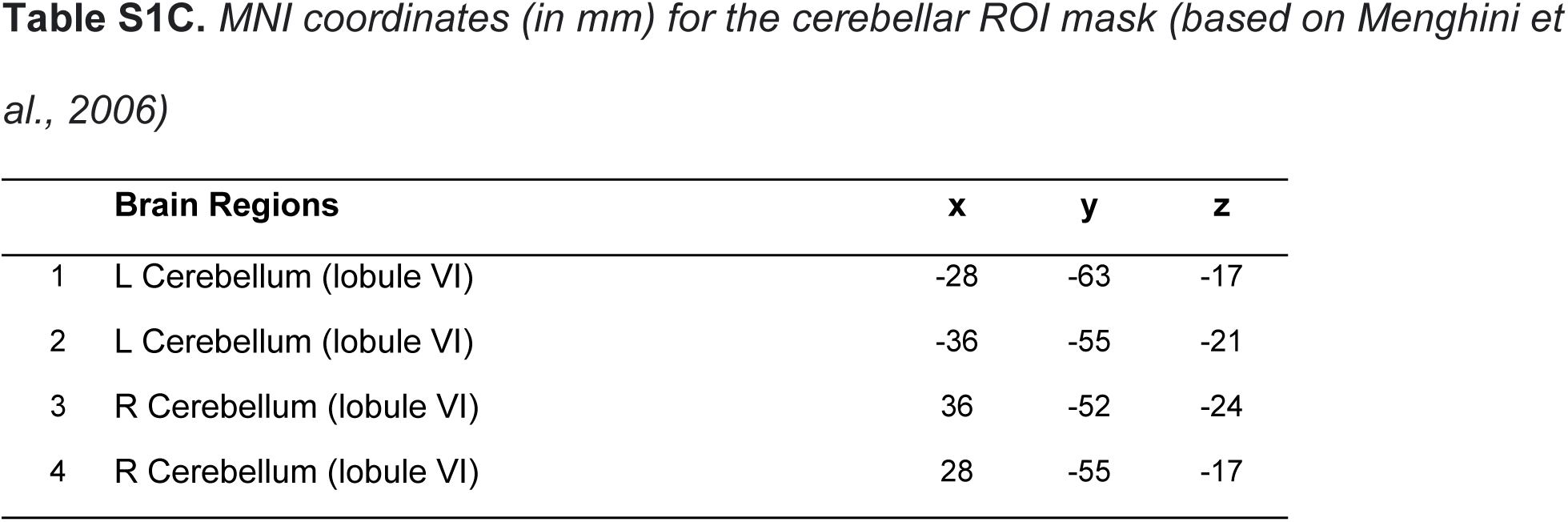
An 8-mm spherical ROI was created with the MarsBaR SPM toolbox (version 0.45; Brett et al., 2002) around each coordinate, taken as the centre of the sphere. L, left; R, right.

**Table S2.** Regions of interest included in the cortical masks for the gray matter volume and cortical thickness analyses.

| Brain Regions |  |
| --- | --- |
| 1 | L Inferior Frontal Gyrus - pars opercularis |
| 2 | L Inferior Frontal Gyrus – pars triangularis |
| 3 | L Superior Frontal Gyrus |
| 4 | L Occipito-temporal Gyrus - Lateral-Fusiform |
| 5 | L Occipito-temporal Gyrus - Medial-Lingual |
| 6 | L Occipito-temporal Gyrus - Medial-Parahippocampus |
| 7 | L Parietal Gyrus - Inferior-Angular |
| 8 | L Parietal Gyrus - Inferior-Supramarginal |
| 9 | L Precentral Gyrus |
| 10 | L Superior-Lateral Temporal Gyrus |
| 11 | L Superior Temporal Gyrus - Planum temporale |
| 12 | R Middle Frontal Gyrus |
| 13 | R Superior Frontal Gyrus |
| 14 | R Occipito-temporal Gyrus - Lateral-Fusiform |
| 15 | R Parietal Gyrus - Inferior-Supramarginal |
| 16 | R Precentral Gyrus |
| 17 | R Superior-Lateral Temporal Gyrus |
| 18 | R Superior Temporal Gyrus - Planum temporale |
| 19 | L Cerebellum |
| 20 | R Cerebellum |

*Note to Table S2.* Between-group differences in cortical regions were assessed using a surface-based approach, while differences in the left and right cerebellum were tested using volumetric analysis (Destrieux et al., 2010). L, left; R, right.

**Table S3.** Tracts of interest included in the white matter mask used for Fractional Anisotropy and Mean Diffusivity analyses (Radwan et al., 2022). L, left; R, right.

| **Brain Regions** |  |
| --- | --- |
| 1 | L Arcuate Fasciculus |
| 2 | L Superior Longitudinal Fasciculus - II dorsal |
| 3 | L Superior Longitudinal Fasciculus - II ventral |
| 4 | L Superior Longitudinal Fasciculus - III |
| 5 | L Inferior Longitudinal Fasciculus |
| 6 | L Inferior Fronto Occipital Fasciculus |
| 7 | L Middle Longitudinal Fasciculus |
| 8 | L Uncinate Fasciculus |
| 9 | L Superior Thalamic Radiation |
| 10 | L Cortico Spinal Tract |
| 11 | R Superior Longitudinal Fasciculus - II dorsal |
| 12 | R Superior Longitudinal Fasciculus - II ventral |
| 13 | R Superior Longitudinal Fasciculus - III |
| 14 | Corpus Callosum – Occipital part |
| 15 | Corpus Callosum – Parietal part |
| 16 | Corpus Callosum – Temporal Part |

**Table S4.**
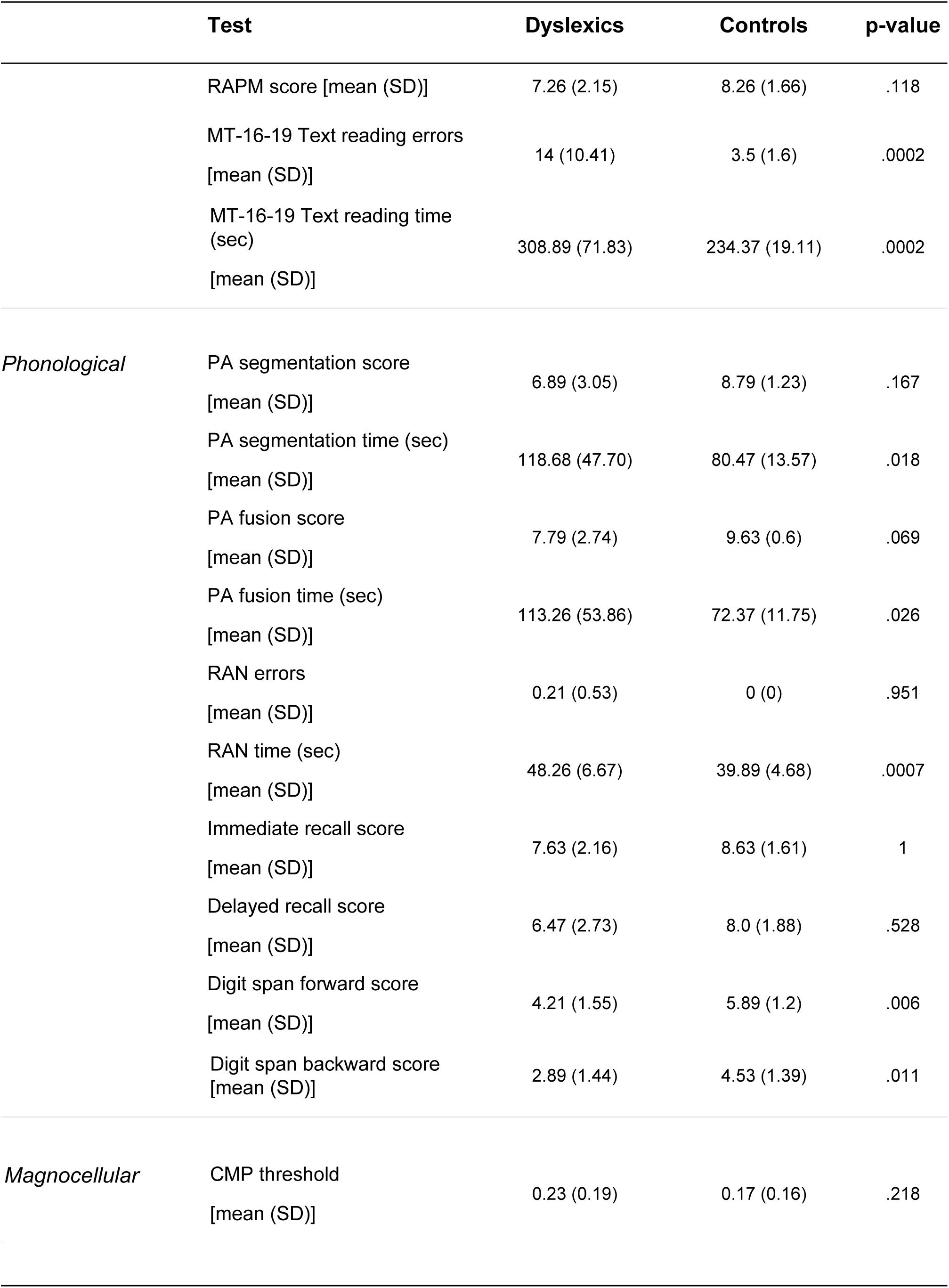
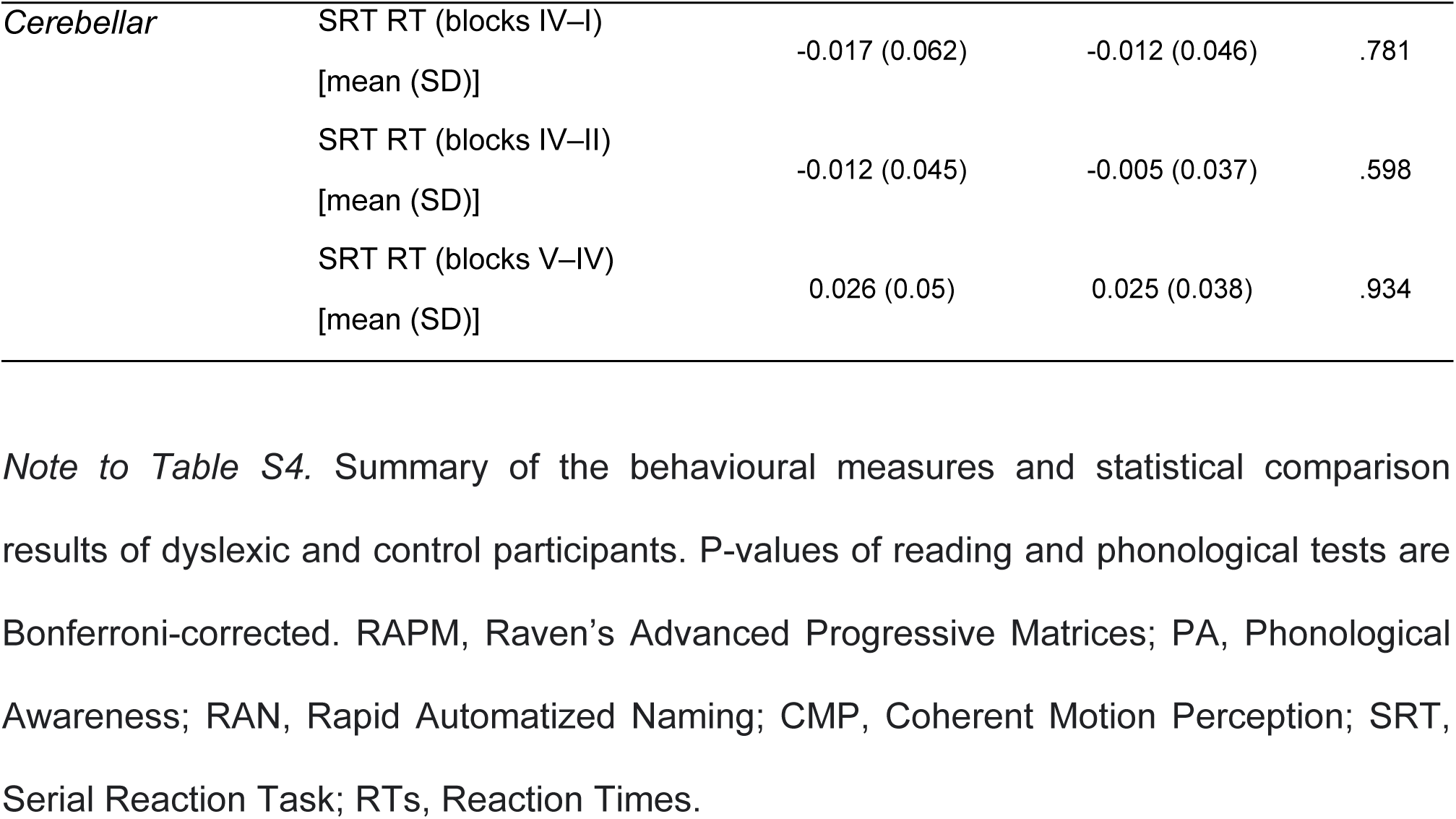
Behavioural results.

|  | Test | Dyslexics | Controls | p-value |
| --- | --- | --- | --- | --- |
|  | RAPM score [mean (SD)] | 7.26 (2.15) | 8.26 (1.66) | .118 |
|  | MT-16-19 Text reading errors<br>[mean (SD)] | 14 (10.41) | 3.5 (1.6) | .0002 |
|  | MT-16-19 Text reading time<br>(sec)<br>[mean (SD)] | 308.89 (71.83) | 234.37 (19.11) | .0002 |
| *Phonological* | PA segmentation score<br>[mean (SD)] | 6.89 (3.05) | 8.79 (1.23) | .167 |
|  | PA segmentation time (sec)<br>[mean (SD)] | 118.68 (47.70) | 80.47 (13.57) | .018 |
|  | PA fusion score<br>[mean (SD)] | 7.79 (2.74) | 9.63 (0.6) | .069 |
|  | PA fusion time (sec)<br>[mean (SD)] | 113.26 (53.86) | 72.37 (11.75) | .026 |
|  | RAN errors<br>[mean (SD)] | 0.21 (0.53) | 0 (0) | .951 |
|  | RAN time (sec)<br>[mean (SD)] | 48.26 (6.67) | 39.89 (4.68) | .0007 |
|  | Immediate recall score<br>[mean (SD)] | 7.63 (2.16) | 8.63 (1.61) | 1 |
|  | Delayed recall score<br>[mean (SD)] | 6.47 (2.73) | 8.0 (1.88) | .528 |
|  | Digit span forward score<br>[mean (SD)] | 4.21 (1.55) | 5.89 (1.2) | .006 |
|  | Digit span backward score<br>[mean (SD)] | 2.89 (1.44) | 4.53 (1.39) | .011 |
| *Magnocellular* | CMP threshold<br>[mean (SD)] | 0.23 (0.19) | 0.17 (0.16) | .218 |
| *Cerebellar* | SRT RT (blocks IV–I) | -0.017 (0.062) | -0.012 (0.046) | .781 |
|  | [mean (SD)] |  |  |  |
|  | SRT RT (blocks IV–II) | -0.012 (0.045) | -0.005 (0.037) | .598 |
|  | [mean (SD)] |  |  |  |
|  | SRT RT (blocks V–IV) | 0.026 (0.05) | 0.025 (0.038) | .934 |
|  | [mean (SD)] |  |  |  |

*Note to Table S4.* Summary of the behavioural measures and statistical comparison results of dyslexic and control participants. P-values of reading and phonological tests are Bonferroni-corrected. RAPM, Raven's Advanced Progressive Matrices; PA, Phonological Awareness; RAN, Rapid Automatized Naming; CMP, Coherent Motion Perception; SRT, Serial Reaction Task; RTs, Reaction Times.

**Table S5.**
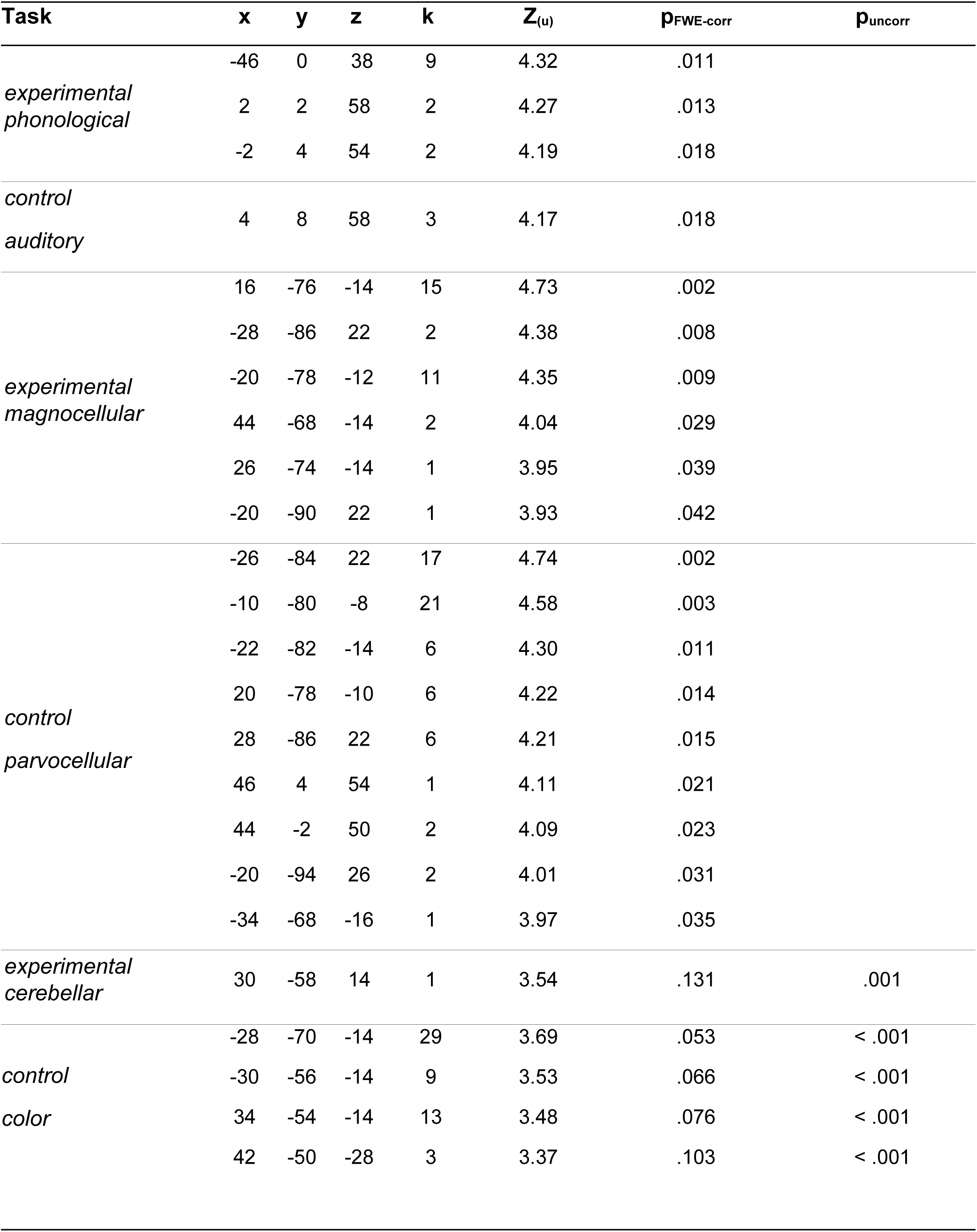
Mismatch response in the whole sample of participants (dyslexics + controls, N = 38)

| **Task** | **x** | **y** | **z** | **k** | **Z<sub>(u)</sub>** | **p<sub>FWE-corr</sub>** | **p<sub>uncorr</sub>** |
| --- | --- | --- | --- | --- | --- | --- | --- |
| *experimental phonological* | -46 | 0 | 38 | 9 | 4.32 | .011 |  |
|  | 2 | 2 | 58 | 2 | 4.27 | .013 |  |
|  | -2 | 4 | 54 | 2 | 4.19 | .018 |  |
| *control auditory* | 4 | 8 | 58 | 3 | 4.17 | .018 |  |
| *experimental magnocellular* | 16 | -76 | -14 | 15 | 4.73 | .002 |  |
|  | -28 | -86 | 22 | 2 | 4.38 | .008 |  |
|  | -20 | -78 | -12 | 11 | 4.35 | .009 |  |
|  | 44 | -68 | -14 | 2 | 4.04 | .029 |  |
|  | 26 | -74 | -14 | 1 | 3.95 | .039 |  |
|  | -20 | -90 | 22 | 1 | 3.93 | .042 |  |
| *control parvocellular* | -26 | -84 | 22 | 17 | 4.74 | .002 |  |
|  | -10 | -80 | -8 | 21 | 4.58 | .003 |  |
|  | -22 | -82 | -14 | 6 | 4.30 | .011 |  |
|  | 20 | -78 | -10 | 6 | 4.22 | .014 |  |
|  | 28 | -86 | 22 | 6 | 4.21 | .015 |  |
|  | 46 | 4 | 54 | 1 | 4.11 | .021 |  |
|  | 44 | -2 | 50 | 2 | 4.09 | .023 |  |
|  | -20 | -94 | 26 | 2 | 4.01 | .031 |  |
|  | -34 | -68 | -16 | 1 | 3.97 | .035 |  |
| *experimental cerebellar* | 30 | -58 | 14 | 1 | 3.54 | .131 | .001 |
| *control color* | -28 | -70 | -14 | 29 | 3.69 | .053 | < .001 |
|  | -30 | -56 | -14 | 9 | 3.53 | .066 | < .001 |
|  | 34 | -54 | -14 | 13 | 3.48 | .076 | < .001 |
|  | 42 | -50 | -28 | 3 | 3.37 | .103 | < .001 |

**Table S6.**
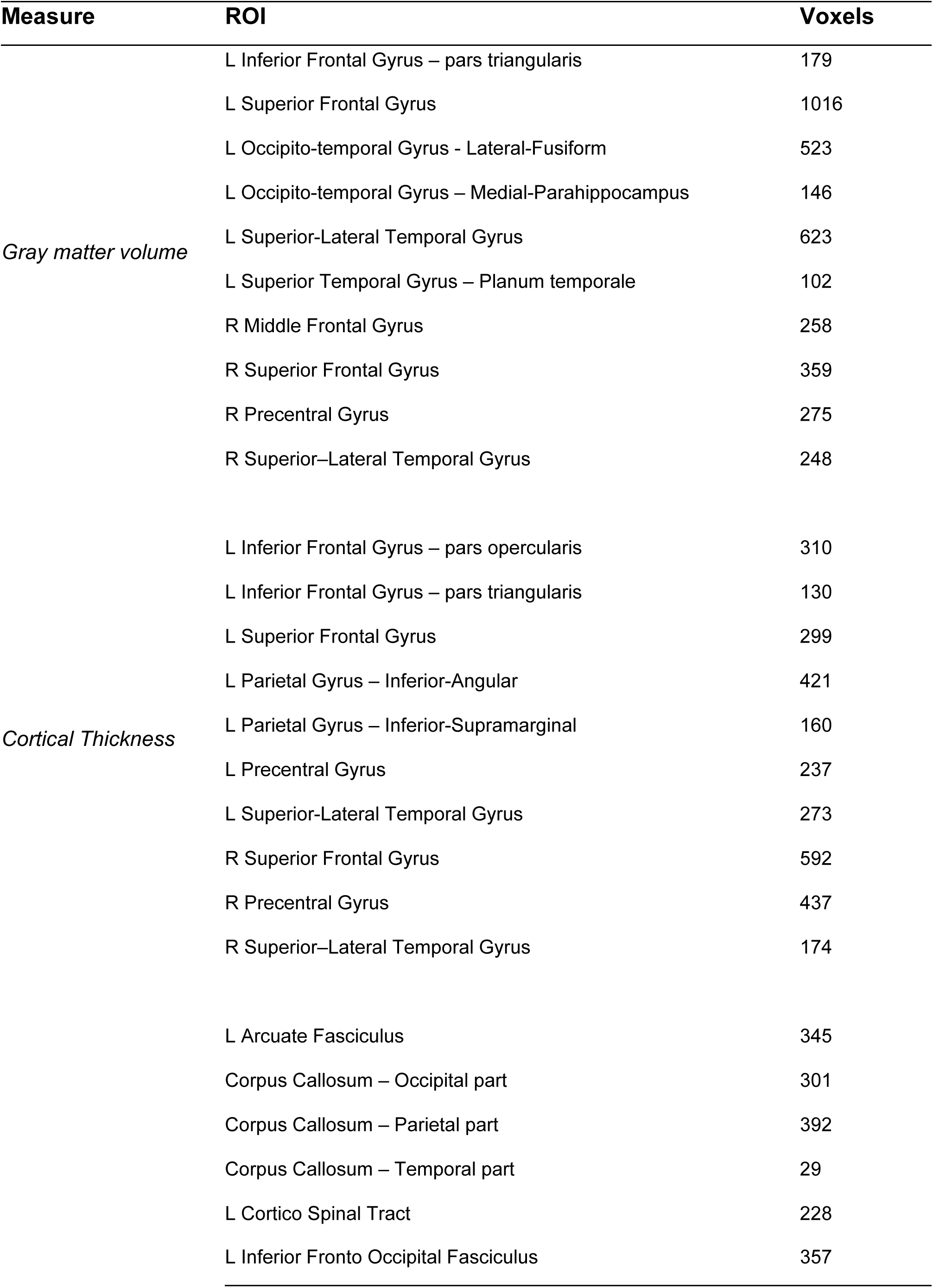
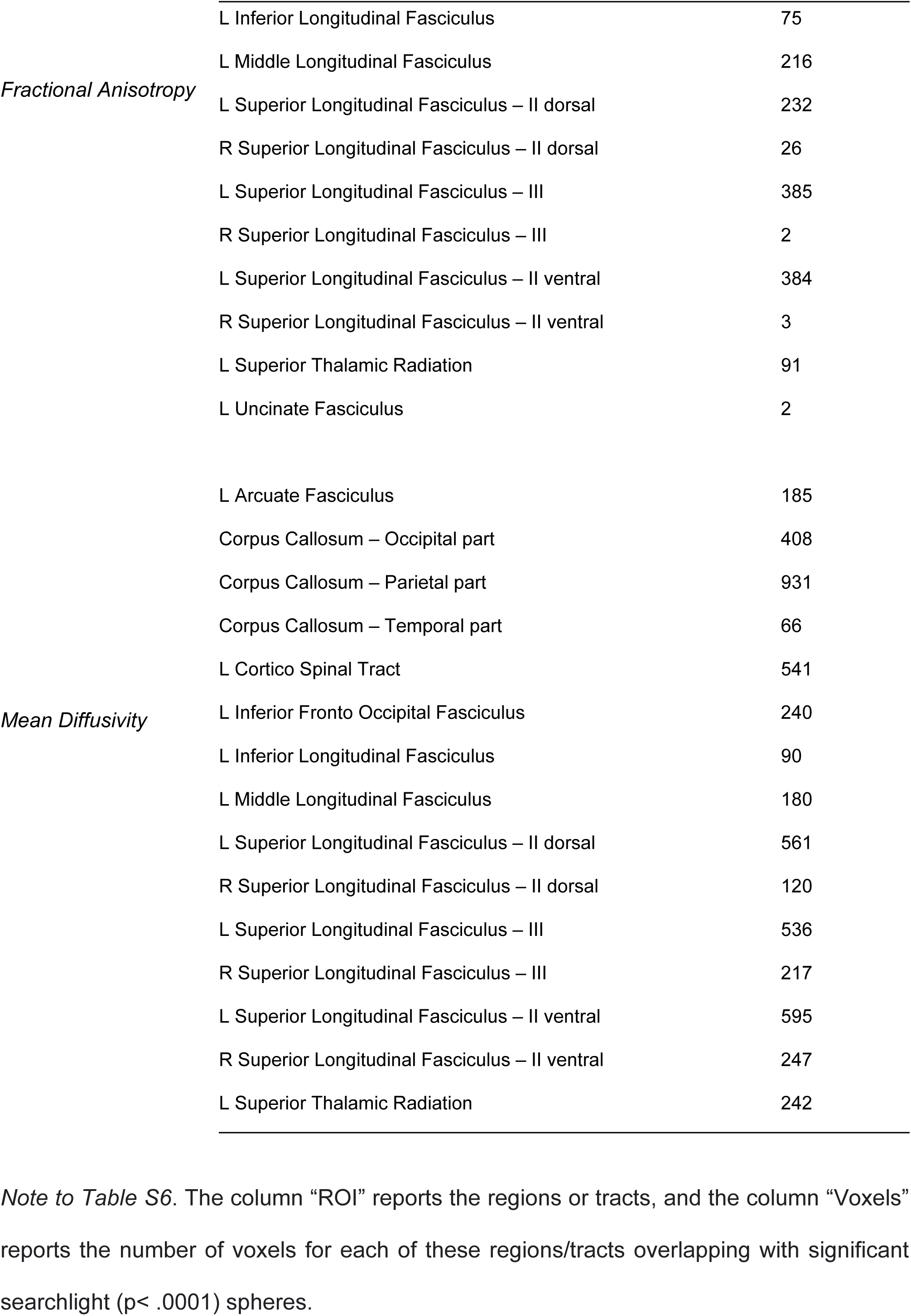
MVPA results.

